# Post-injury hydraulic fracturing drives fissure formation in the zebrafish basal epidermal cell layer

**DOI:** 10.1101/2022.05.21.492930

**Authors:** Andrew S. Kennard, Mugdha Sathe, Ellen C. Labuz, Christopher K. Prinz, Julie A. Theriot

**Author notes:** Contributed equally (Andrew S. Kennard, Mugdha Sathe). Department of Biology, University of Massachusetts, Amherst, MA, USA.

## Abstract

The skin epithelium acts as the barrier between an organism’s internal and external environments. In zebrafish and other freshwater organisms, this barrier function requires withstanding a large osmotic pressure differential. Wounds breach this epithelium, causing a large disruption to the tissue microenvironment due to the mixing of isotonic interstitial fluid with the external hypotonic fresh water. Here we show that, following acute injury, the larval zebrafish epidermis undergoes a dramatic fissuring process that resembles hydraulic fracturing, driven by the influx of external fluid. The fissuring starts in the basal epidermal layer nearest to the wound, and then propagates at a constant rate through the tissue spanning over one hundred micrometers; during this process the outermost superficial epidermal layer remains intact. Fissuring is completely inhibited when larvae are wounded in an isotonic external media, suggesting that osmotic pressure gradients drive fissure. Additionally, fissuring partially depends on myosin II activity, as its inhibition reduces fissure propagation away from the wound. During and after fissuring, the basal layer forms large macropinosomes (with cross-sectional areas ranging from 1-10 µm^2^), presumably to clear the excess fluid. We conclude that excess external fluid entry through the wound and subsequent closure of the wound through actomyosin purse string contraction in the superficial cell layer causes fluid pressure buildup in the extracellular space of the zebrafish epidermis. This excess fluid pressure causes tissue to fissure, and eventually the fluid is cleared through macropinocytosis.

## Introduction

A key function of many animal epithelial tissues is to establish and maintain extreme asymmetries in environmental composition between their apical and basal surfaces. For example, the mammalian gastric epithelium establishes a pH gradient between the stomach lumen (pH ∼3) and the rest of the body (pH ∼7.4) (Schreiber et al., 2004), frog epidermis generates an electrical charge imbalance leading to a trans-epithelial electrical potential of tens of millivolts (Ferreira et al., 2016; Robinson, 1983), and larvae of freshwater zebrafish withstand over 20-fold differences in osmolarity across their epidermis (Kennard and Theriot, 2020; Krens et al., 2017). Epithelial injury disrupts such gradients, and wound healing is critical for epithelia to re-establish these gradients by re-sealing injured tissue and preventing unregulated transepithelial flux.

Wound healing is a complex process that includes different stages depending on the animal species and the nature of the wound. A common and prominent phase of wound healing is reepithelialization, a collective behavior in which epithelial cells detect a nearby injury and migrate in a coordinated fashion towards the wound, closing off the damaged area (Richardson et al., 2016). Reepithelialization is driven by actomyosin-based processes, including the development of supracellular actomyosin purse-string cables that tighten to contract the wound margin, and the directed actin-based migration of epithelial cells towards the site of injury (Abreu-Blanco et al., 2012; Eming et al., 2014; Martin and Lewis, 1992; Richardson et al., 2016; Rothenberg and Fernandez-Gonzalez, 2019). Physical coordination of tissue connectivity through cell-cell junctions is also critical for reepithelialization, and the level of adhesion between cells is carefully regulated to facilitate movement while retaining tissue organization (Hunter et al., 2015; Nunan et al., 2015; Tetley et al., 2019). For example, during healing of mouse skin wounds, cells downregulate tight junctions and E-cadherin-based adherens junctions while retaining desmosomal junctions, a process that is believed to loosen the tissue and facilitate migration towards the wound (Nunan et al., 2015).

Much remains unresolved concerning how disruptions to transepithelial gradients following epithelial injury affect reepithelialization and wound healing. These mechanisms may be especially important in the epidermis, which generates extreme gradients in ionic electrical potential and fluid composition between interstitial fluid and the outside world (Barker et al., 1982). In some contexts, environmental changes resulting from gradient disruption have been shown to have a signaling function, serving as cues of tissue damage (Enyedi and Niethammer, 2015). For example, short-circuiting of trans-epithelial electrical potentials can create electric fields within the epidermis that guide cell migration and tissue regeneration (Ferreira et al., 2016; Kennard and Theriot, 2020; Kucerova et al., 2011; Reid et al., 2005; Sun et al., 2011), and nuclear swelling driven by osmotic shock can activate signal transduction cascades that drive reepithelialization and immune cell infiltration of damaged tissue (Chen et al., 2019; Enyedi et al., 2016, 2013; Gault et al., 2014). Beyond activating signal transduction programs, it also seems plausible that gradient disruption could provide a powerful driving force for physical processes, such as fluid flow, which could affect the structure or organization of epithelial tissue during wound healing.

We were particularly interested in how transepithelial gradient disruption would affect the structure of healing epithelial tissue. Zebrafish are an ideal system for investigating the relationship between gradient disruption and wound healing, due to their power as a model organism and their aqueous, freshwater lifestyle. The excellent optical properties of larval zebrafish epidermis and the availability of genetic tools for live fluorescence imaging have made zebrafish larvae a prominent model system for understanding the rapid mechanisms of wound repair in the first few minutes after injury (Enyedi et al., 2013; Franco et al., 2019; Gault et al., 2014; Kennard and Theriot, 2020; Poplimont et al., 2020; Yoo et al., 2012). The zebrafish larval epidermis is composed of two layers of cells: a superficial layer that maintains barrier function via tight junctions, and a basal layer of proliferative cells that can migrate on the basement membrane/extracellular matrix (ECM) and can perform other tissue maintenance functions, including phagocytosis of apoptotic cells (Arora et al., 2020; Rasmussen et al., 2015; Sonawane et al., 2005) (**Figure 1A-B**). Previous work has shown that the two layers respond to injury using distinct wound closure mechanisms: superficial cells assemble an actomyosin purse string around the wound margin, while basal cells actively migrate towards the wound using actin-rich lamellipodia (Franco et al., 2019; Gault et al., 2014). Crucially, zebrafish are freshwater organisms capable of withstanding more than 20-fold osmotic gradients between interstitial fluid and the surrounding environment (Krens et al., 2017). Several environmental cues contribute to wound detection in zebrafish, including the aforementioned osmolarity-induced cell swelling and the establishment of electric fields directed towards the wound (Enyedi et al., 2016; Gault et al., 2014; Kennard and Theriot, 2020). While effects of these rapid, injury-induced changes in the cellular environment have been identified at the earliest stages of wound healing, it is not known how these substantial environmental disruptions affect later stages of wound healing, or even how the pre-wounding interstitial environment is restored.

**Figure 1.**
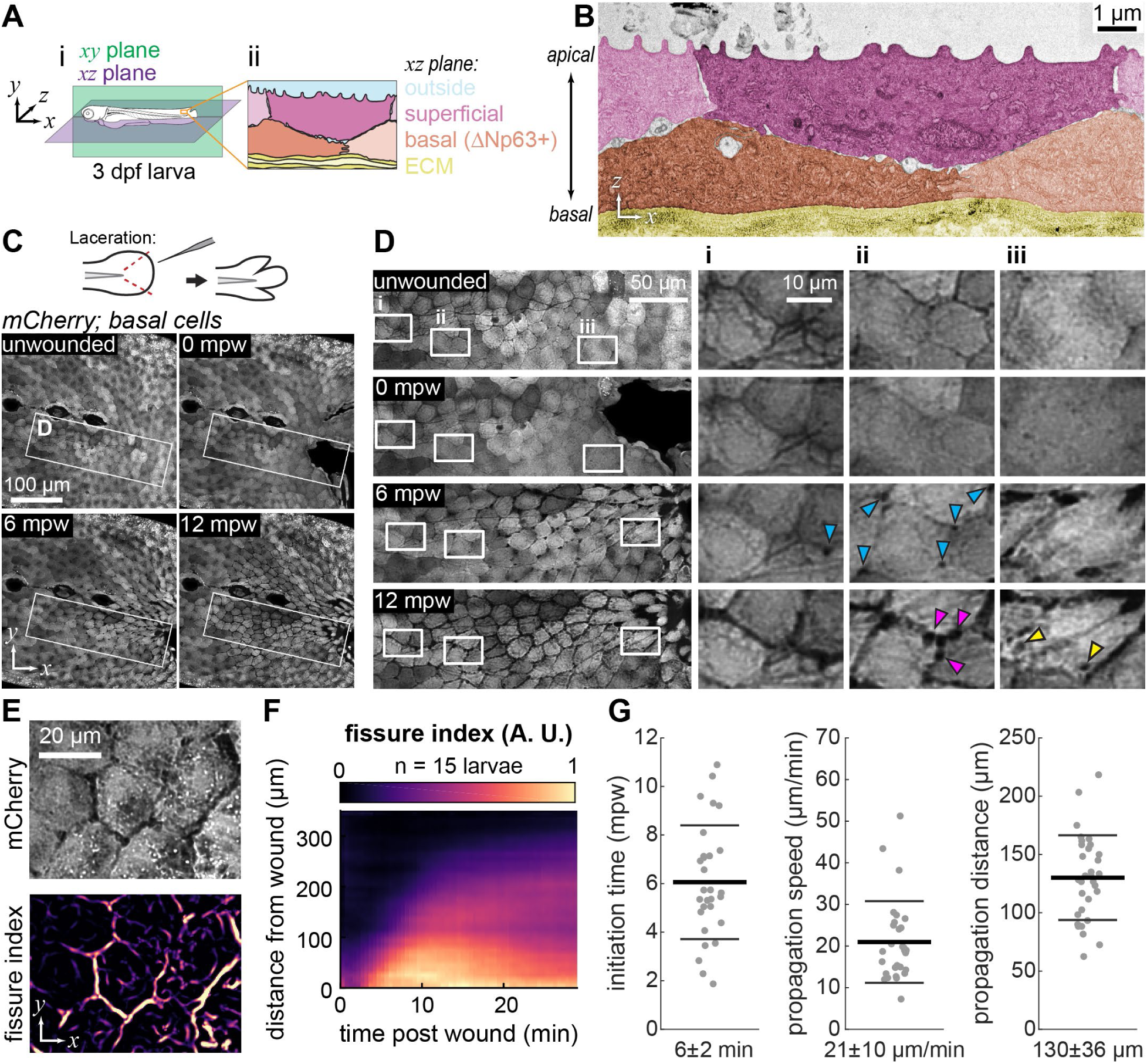
Fissures propagate between epidermal cells during wound healing. **(A)** (i) Schematic of a 3 dpf zebrafish larva defining the *xy* (sagittal) and *xz* (coronal) planes used throughout the paper. (ii) *xz* section through the larval epidermis shows the bilayered structure, formed of superficial layer and the basal layer (specified by the *ΔNp63* promoter) resting upon the extracellular matrix (ECM). **(B)** Electron micrograph of unwounded 3 dpf epidermis, pseudo-colored according to (Aii). **(C)** Maximum intensity projections of 3 dpf larva expressing cytoplasmic mCherry in basal cells *(TgBAC(*Δ*Np63:Gal4); Tg(UAS:mCherry))* at different time points after wounding. Schematic of the laceration procedure is illustrated at the top. **(D)** Insets from (C) revealing the propagation of fissures between basal cells over time after wounding. Blue arrowheads indicate the appearance of dark puncta at tricellular junctions that precedes full fissuring. Yellow arrowheads indicate large dark puncta that appear within cells after fissuring has occurred. Magenta arrowheads highlight the irregular, “beads-on-a-string” morphology that fissures develop over time. Insets i-iii follow specific cells over time as they migrate to the wound. The cell followed in inset iii migrates the most, consistent with its position nearest the wound. **(E)** Example of the fissure index image computed from the basal cell mCherry signal. **(F)** Kymograph of average fissure index demonstrating initial linear propagation of fissuring away from the wound followed by slower propagation after about 12 minutes. Fissure index is averaged over 15 larvae. **(G)** Quantified fissure dynamics, including the initiation time of fissuring, the propagation speed during the initial linear propagation interval, and the spatial extent of linear propagation. Each dot represents an individual larva (n ≥ 29), and the thick and thin bars represent the mean +/- 1 standard deviation.

Studying reepithelialization in larval zebrafish epidermis, we report that—in addition to contributing to wound detection—the influx of external media at the wound site also permeates deep into the epidermis through fissures that emerge between basal epidermal cells after injury. These fissures propagate at constant speed away from the site of injury for over 100 µm— roughly 10 cell diameters—and fissuring is promoted both by osmotic pressure gradients and by myosin-based tissue contractility. After fissuring, cells remain connected via tethers that contain adherens junctions and desmosomes, and they uptake the surplus of externally derived fluid through an increase in macropinocytosis. Our data show that the rapid influx of excess extracellular fluid inevitably associated with a breach in the epidermis causes a previously unanticipated set of events, namely fissuring in the epithelial tissue, that is eventually cleared by macropinocytosis. These results emphasize the critical contribution of large-scale physical processes such as fluid flow in shaping the wound response in zebrafish larvae.

## Results

### Fissures emerge between basal epidermal cells after they have stopped migrating towards a wound

Using a previously developed laceration wounding technique (Kennard et al., 2021), we sought to identify the sequence of events occurring towards the end of reepithelialization, as cells slow down and re-form a stationary epithelium. In this wounding approach, a glass needle is used to impale and tear the posterior end of the tailfin, generating two full-thickness cuts just posterior to the notochord. We have found that this wounding approach elicits a robust migratory wound response beginning almost immediately after injury and persisting for about 15 minutes, during which time basal cells form actin-rich lamellipodial protrusions and migrate toward the injured region (Kennard and Theriot, 2020). Surprisingly, we found that, as basal epidermal cells slowed down, they began to separate from each other, so that gaps several micrometers wide appeared between cells, which we termed “fissures” (**Figure 1C-D, Video S1**). Fissures were most easily observed in larvae in which basal epidermal cells expressed a cytoplasmic fluorescent protein (mCherry) as a volume marker. This fissuring was strikingly regular, and appeared to propagate in a wave through the tissue, beginning at the wound site and emanating anteriorly (**Figure 1D**). The rate at which fissures propagated anteriorly did not appear to vary strongly with position along the dorsal-ventral axis, although fissuring frequently initiated at the midline. Fissures tended to decrease in width with increasing distance from the wound, and fissures thinned over time (**Video S1**), although they persisted for at least 3 hours after wounding (**Figure S1**).

Upon closer inspection, it became clear that small gaps opening at tricellular junctions preceded fissure formation between adjacent cells (**Figure 1D**, blue arrowheads). Following fissure formation, cells began to accumulate numerous dark puncta internally, representing decreases in the local fluorescence intensity of the cytoplasmic fluorescent protein signal, as would be expected for fluid-filled endosomes or macropinosomes (**Figure 1D**, yellow arrowheads). These dark puncta varied in size but sometimes reached several micrometers in diameter. In tandem with puncta accumulation, the fissures on the periphery of each cell transitioned from smooth, uniform gaps with a constant width to an uneven “beads-on-a-string” morphology, with the width of the gap varying along each cell-cell junction (**Figure 1D**, magenta arrowheads).

To quantitatively compare the dynamics of fissuring across many larvae and conditions, we developed a computational pipeline to identify separated cell boundaries in each frame of time-lapse movies (see Methods). To emphasize fissures, this pipeline used a ridge detection filter, which produces an image in which the intensity value of each pixel is related to the degree to which the neighborhood of that pixel resembles a dark linear structure in the original image; we termed this measurement the “fissure index” (**Figure 1E, Figure S2A**). To reveal the extent and speed of fissure propagation through the tissue, we binned the average value of the fissure index as a function of distance from the wound and time after wounding, producing a fissure kymograph, which reports on areas with significant ridge detection over time (**Figure 1F**). Fissure kymographs averaged from multiple larvae confirmed our observation that fissures first propagate at a constant speed through the tissue, followed by a phase of slower propagation. (**Figure 1F**).

To extract the initiation time, speed, and distance of fissure propagation, we fit a line to the linear propagation portion of the kymograph (**Figure S2B**). This analysis showed that fissures initiate 6 ± 2 minutes post wounding (mpw) (s.d., *n* = 29 larvae) and propagate through the tissue at 21 ± 10 µm/min to a distance of 130 ± 36 µm (**Figure 1G**). Previously, we had shown that, during the migratory phase of the wound response in zebrafish basal epidermis, cell migration initiates in an anteriorly propagating wave, analogous to fissure formation (Kennard and Theriot, 2020). Intriguingly, the spatial extent and propagation speed of this wave of cell movement (200 µm and 40 µm/min) is comparable to the extent and speed of fissure propagation (130 µm and 21 µm/min), while the initiation of the wave of cell movement begins almost immediately after injury, preceding fissure propagation. Overall, our quantification of fissure dynamics reveals that basal cell fissures propagate away from the wound at a constant rate for many cell diameters before slowing down.

### Separated basal cells remain connected by thin tethers, while the superficial layer remains intact

Given the importance of cell-cell adhesion for many collective tissue behaviors including migration of epithelial monolayers (Friedl and Mayor, 2017), the emergence of fissures between basal cells was unexpected. To interrogate the structure of fissures in more detail, we performed thin-section transmission electron microscopy on unwounded larvae and larvae fixed at 2, 7, and 20 mpw. As previously shown (Le Guellec et al., 2004; O’Brien et al., 2012; Sonawane et al., 2005), basal cells and superficial cells of unwounded larvae were tightly apposed in an interlocking pattern: the cross-section of each cell was tallest in the middle above the nucleus and narrowed toward the cell-cell junctions at the edges (**Figure 1B**, **Figure 2A**).

**Figure 2.**
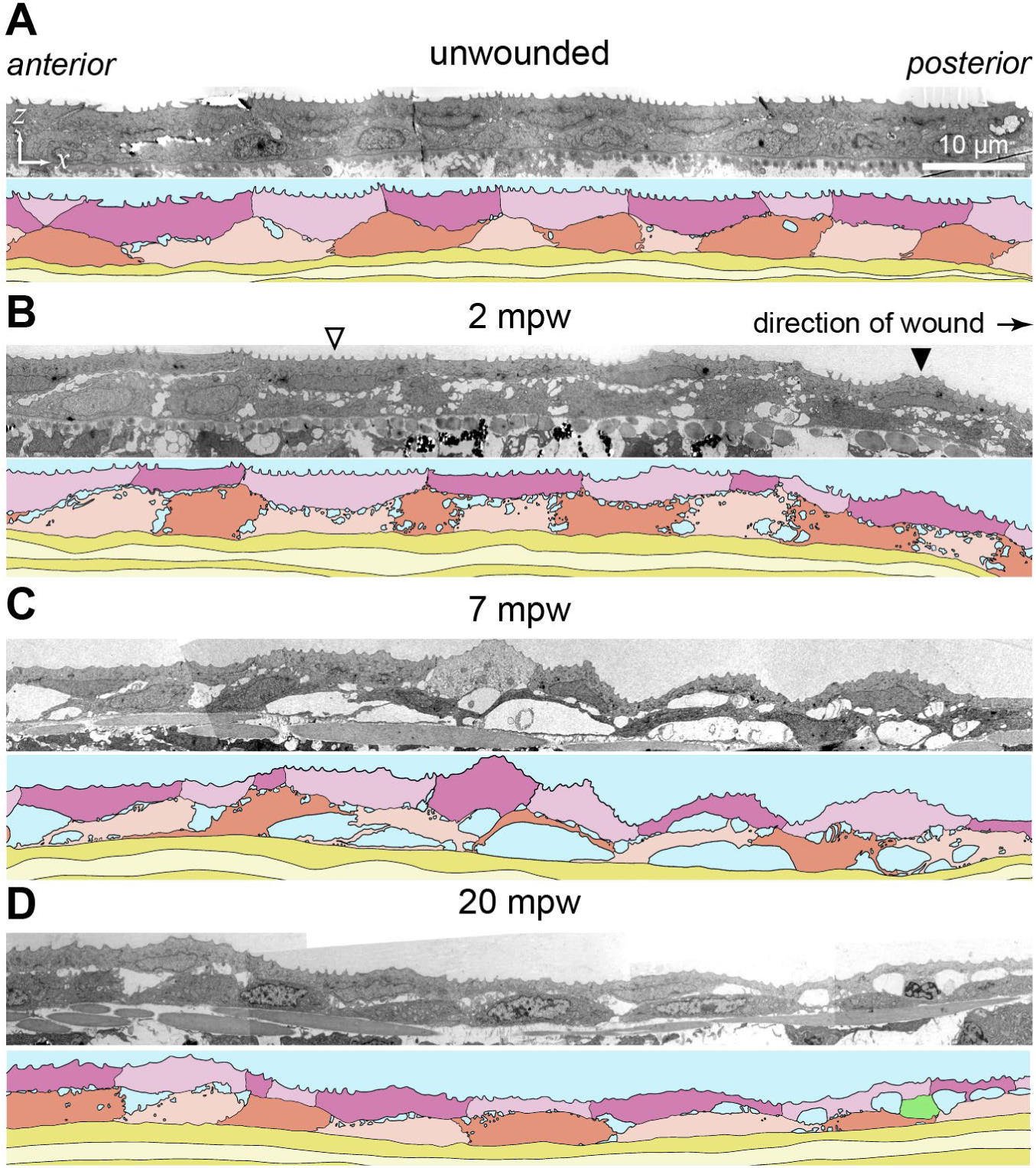
Gaps between basal cells develop soon after wounding. **(A-D)** Electron micrograph *xz* planes of 3 dpf epidermis before wounding (A) or at various times after wounding (B-D). Each micrograph is accompanied by a camera lucida drawing to guide the eye. Coloring of camera lucida drawings matches that in Figure 1A; the cell in green in panel (D) is an immune cell. Gaps between cells emerge as early as 2 mpw, coalesce by 7 mpw, and by 20 mpw they appear at cell-cell junctions at the interface between superficial and basal cells. White arrowhead in (B): superficial cell further away from the wound retains the flat apical surface and curved basal surface seen in unwounded cells. Black arrowhead in (B): superficial cell closer to the wound has an inverted shape, with a curved apical surface and a flattened basal surface.

This characteristic, interlocking three-dimensional epidermal architecture is in line with the geometry previously established for frog larval epidermis (Savost’yanov and Grefner, 1998). In contrast to the well-connected, unwounded epidermis, many small empty spaces were visible between basal cells and at the basal-superficial interface in epidermal cross-sections 2 mpw (**Figure 2B**). Manual segmentation of cells and the gaps between them indicates that gaps make up ∼12% of the total epidermal cross-sectional area at 2 mpw, an increase from ∼2% in unwounded tissue. While the cross-sections of superficial cells retained their unwounded shape further from the wound (**Figure 2B**, white arrowhead), closer to the wound superficial cells were no longer interlocking with basal cells (**Figure 2B**, black arrowhead); instead, superficial cells bulged on the apical side and were flatter on the basal side.

At 7 mpw, when fissures have typically just appeared (as judged by live animal confocal imaging), basal cells were highly elongated along the direction of the wound, consistent with migration typically occurring at that time (Gault et al., 2014; Kennard and Theriot, 2020). At this timepoint the gaps between basal cells were much larger than at 2 mpw, especially between the basal cells and the basement membrane (**Figure 2C**). At 20 mpw, when migration has ceased and fissures are still readily observable by light microscopy, basal cells had adopted a flattened ovoid shape and large, rounded gaps were observed at the junctions between adjacent basal cells (**Figure 2D**). These were morphologically distinct from the small gaps observed at 2 mpw. Despite these large gaps, basal cells still frequently maintained lateral contacts with their neighbors. At all timepoints, superficial cells retained close lateral contacts, suggesting that fissures did not occur in this layer. Additionally, gaps were readily seen both at the lateral interfaces between basal cells, and at the interfaces between basal and superficial cells, suggesting that fissures form around all basal cell surfaces, although they are only detectable at lateral surfaces in maximum-intensity projections of confocal images. To rule out fixation artifacts, we observed wound closure in larvae expressing LifeAct in basal cells and observed the effect on fissure and overall tissue structure during fixation with our EM fixative. We found that the shape and organization of fissures that was appreciable with light microscopy was not altered by fixation (**Figure S3**). Overall, our EM observations indicate that gaps between basal cells occur rapidly after wounding—even before such gaps are detectable by light microscopy— and that over time they coalesce, while the superficial layer remains intact throughout the wound healing process.

Electron microscopy revealed that the fissuring we observed in light microscopy did not completely eliminate the contacts between cells (**Figure 2**). To test whether E-cadherin was present at these persistent contacts, we wounded larvae which expressed mCherry in the basal layer and zebrafish E-cadherin (Cdh1) tagged with sfGFP in both superficial and basal layers (Yamaguchi et al., 2019). Using the basal-cell specific mCherry, we computationally distinguished the E-cadherin signal arising from basal or superficial layers, and compared the dynamics of E-cadherin in each layer to the dynamics of fissures, revealed by dark gaps in the mCherry channel. We observed that E-cadherin was maintained at basal and superficial junctions throughout the wound-healing process (**Figure 3A, Video S2**). The retention of E-cadherin at junctions despite the reduction in cytoplasmic signal there is consistent with the many membrane tethers observed in electron microscopy (**Figure 2**), which would have a high surface area to volume ratio and thus preferentially enrich for membrane-bound markers like E-cadherin over cytoplasmic markers. Although E-cadherin remained localized at junctions throughout wound healing, it is important to note that the intensity of E-cadherin signal in both basal and superficial layers decreased markedly during the course of our experiments; due to the low level of expression from a bacterial artificial chromosome (BAC)-based native *cdh1* promoter and the relatively high bleaching rate of sfGFP, it was not possible to distinguish between sfGFP photobleaching and possible E-cadherin degradation. We also looked at cell-cell junctions at high magnification in our electron microscopy samples. We saw that the tethers connecting cells frequently terminated in an electron-dense plaque (**Figure 3B**, blue arrowhead) reminiscent of desmosomal plaques, with many intermediate filaments extended from the plaques (**Figure 3B**, yellow arrowhead). Taken together, our confocal and electron microscopy of cell-cell junctions suggests that adherens junctions and desmosomes may remain partially intact during fissuring.

**Figure 3.**
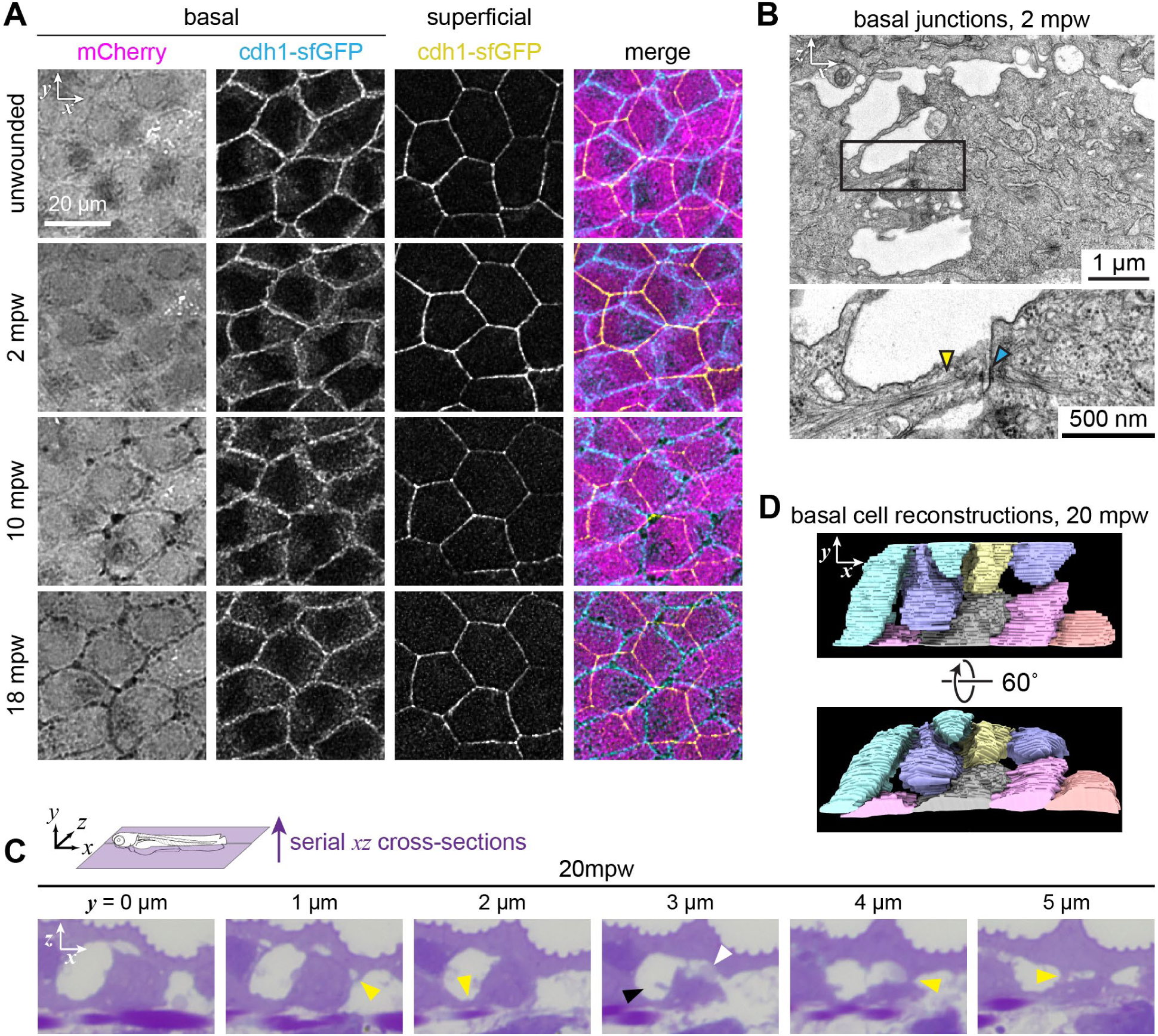
Fissured cells remain connected through cell-cell junctions. **(A)** Surface projections (see Methods) from a timelapse following wounding of a 3 dpf larvae expressing mCherry in basal cells and E-cadherin in basal and superficial cells *(TgBAC(*Δ*Np63:Gal4); Tg(UAS:mCherry); TgBAC(cdh1:cdh1-sfGFP))*. E-cadherin remains localized at the cell periphery in both layers, despite fissuring in the basal layer. To facilitate comparison of the distribution of E-cadherin, image intensity was bleach-corrected at each timepoint by rescaling (see Methods). **(B)** Electron micrograph of the cell-cell junction between two basal cells at 2 mpw. Blue arrowhead in the inset indicates the electron-dense staining characteristic of desmosomal plaques, and the yellow arrowhead indicates intermediate filaments. **(C)** Serial histological sections of epidermis at 20 mpw, taken at 1 µm increments along the *y* (dorsal-ventral) axis. White arrowhead: the gaps on either side of the basal cells are revealed in the *y* = 3 µm section to be contiguous across the apical surface of the cell. Black arrowhead: large rounded gap between two basal cells. Yellow arrowheads: tethers connect basal cells to other basal cells and to superficial cells. **(D)** 3D reconstruction of basal cells from 20 mpw larva shown in (C). Gaps between cells resemble fissures in confocal microscopy and are contiguous with each other along the interface between basal and superficial layers.

To reconcile the appearance of fissures between cells in confocal microscopy with the persistence of tethers and adhesions between cells, we sought to determine the structure of fissures by imaging larger volumes of tissue than were practical with electron microscopy. By cutting semi-thin 0.5 µm sections from resin-embedded samples of 20 mpw larvae, we obtained 20 µm of serial cross-sections for histological imaging and three-dimensional tissue reconstruction (**Video S3**). This was enough to observe the arrangement of fissures around multiple cells. Consistent with our electron microscopy observations, this histological imaging emphasized that fissures appear at both the lateral and apical surfaces of basal cells (**Figure 3C**, black and white arrowheads, respectively), while only lateral fissures are easily detected in confocal microscopy. We also observed many tethers in our histology imaging, typically near the basement membrane or between superficial and basal cells (**Figure 3C**, yellow arrowheads). Importantly, 3D reconstructions of the serial-section histology revealed the continuity of fissures across lateral and apical surfaces (**Figure 3D**). Thus, fissures form an interconnected network at the interface between basal and superficial cell layers, while attachments between cells in both layers persist via membrane tethers.

### Fissures are filled with externally-derived fluid and form via wound contractility and osmotic pressure

The interconnected, channel-like structure of epidermal fissures was evocative of hydraulic fracturing, or “fracking,” which has previously been observed in epithelial monolayers and during early embryogenesis (Casares et al., 2015; Dumortier et al., 2019). We therefore hypothesized that fissuring could be caused by the infiltration of external fluid into the epidermis upon tissue injury. To test whether fissures contained external fluid, we pulsed 10 kDa fluorescently labeled dextran into the external medium surrounding wounded larvae for 7 minutes and imaged the epidermis after washing out excess dextran from the medium (which took roughly 3 minutes, see Methods). We observed that fluorescent dextran readily entered the wounded tissue, and anterior propagation of the fluorescent signal exactly coincided with fissure formation (**Figure 4A** and **Video S4**). Intriguingly, in addition to colocalization of fluorescent dextran with fissures at lateral cell-cell junctions, dextran was also observed in channels crossing the middle of the cell (**Figure 4A**, white arrowheads, see inset), consistent with the fissures at the superficial-basal layer boundary observed in electron microscopy and histology (**Figure 3**). This experiment demonstrates that epidermal fissures contain external fluid.

**Figure 4.**
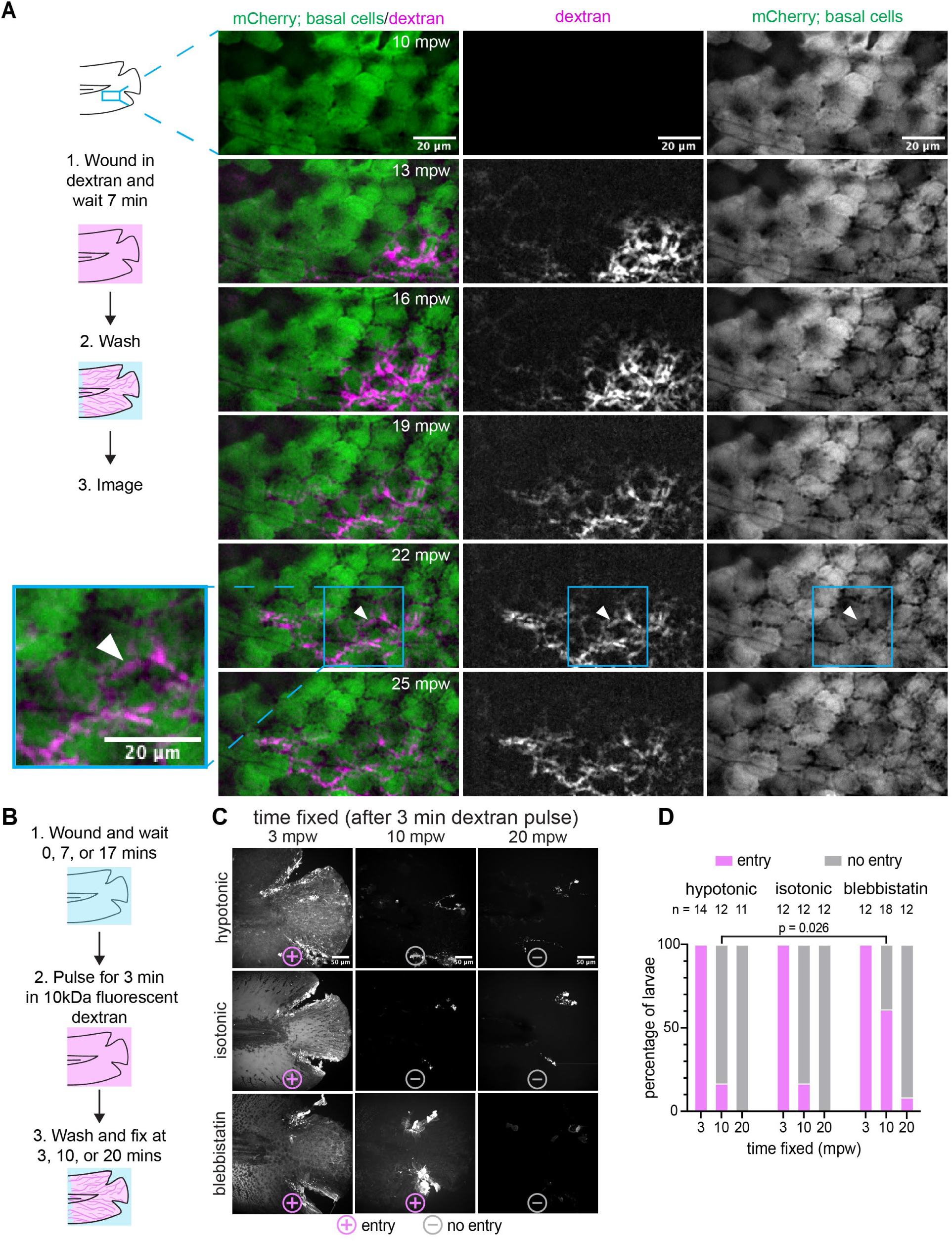
External dextran enters via the open wound *in vivo* and in fixed larvae. **(A)** (Left) Schematic of the experimental workflow. Briefly, the larva was wounded in the presence of TMR-Dextran and incubated for 7 minutes. Larva was then washed with E3 media for roughly 3 minutes and was imaged live with 1 minute intervals. White arrowhead at 22 mpw illustrates an example of dextran flow across the middle of a cell rather than along a cell boundary, possibly due to flow across the apical surface of the cell. (Right) Maximum intensity projections from a timelapse following wounding of a 3 dpf larva expressing mCherry in the basal cells (*TgBAC(*Δ*Np63:Gal4); Tg(UAS:mCherry)*) that were wounded in the presence of E3 media supplemented with 2 mg/ml 10 kDa TMR dextran. **(B)** Visual diagram of the workflow for fixed dextran assays. (1) Larva were wounded in either E3 media (hypotonic), 270 mM sorbitol (isotonic medium) or E3 media supplemented with 50 µM para-nitro blebbistatin. (2) After 0, 7, and 17 minutes of waiting, the larvae were with pulsed with 2mg/ml of TMR-dextran (hypotonic and isotonic) or AF680-dextran (blebbistatin) for 3 minutes. (3) Larvae were washed, fixed and imaged. **(C)** Maximum-intensity projections from the fixed larvae processed as described in (B). Larvae were scored +/– for dextran entry, depicted with symbols below each image **(D)** Percentage of larvae in each condition scored for dextran entry. The number of larvae for each condition is shown above the corresponding bar. A significantly higher proportion of larvae were infiltrated with dextran after 10 mpw in blebbistatin vs. hypotonic medium (*p* = 0.026, Fisher’s exact test).

Hydraulic fracturing of a tissue can be driven by a large increase in hydrostatic pressure of the interstitial fluid. We reasoned that the interstitial fluid could only become pressurized after the wound was closed, as an open wound allows fluid to flow out of the tissue into the extraembryonic environment, rather than stay in the tissue and become pressurized. To determine when the epidermal barrier function recovered after wounding, we applied 3 minute pulses of fixable fluorescent dextran to wounded larvae at different times post wounding, fixed larvae immediately after the pulse, then imaged them to look for the presence of dextran within the tissue (**Figure 4B,C**). In control conditions, dextran was observed within all larvae that were wounded in the presence of fluorescent dextran and then fixed at 3 mpw. However, only 17% of larvae pulsed with fluorescent dextran starting at 7 mpw and then fixed at 10 mpw showed significant fluorescent signal within the tissue. Finally, no larvae exposed to the pulse of fluorescent dextran starting at 17 mpw and fixed at 20 mpw exhibited internal fluorescence (*n* > 11 larvae per time point) (**Figure 4D**). This suggested that the majority of wounds are largely sealed at or before 7 mpw, coinciding with the typical time at which fissure formation and propagation are readily observable by light microscopy (**Figure 1G**).

The pressure to drive tissue fracking could arise from osmotic pressure gradients, which would drive water entry from the hypotonic external medium into the tissue. Additionally, contractile forces at the wound margin could force fluid into the tissue, particularly after the wound has sealed to prevent backflow of fluid into the external environment. To test the contribution of tissue contractility to the infiltration of external fluid, we inhibited tissue contractility with the myosin II inhibitor blebbistatin. To test the contribution of osmotic pressure gradients, we adjusted the external osmolarity with sorbitol to match the osmolarity of internal fluid without disrupting osmolarity-independent ionic or electrical wound cues (Kennard and Theriot, 2020; Krens et al., 2017). We first determined the effect of these perturbations on the ability of the epidermis to reestablish barrier function after injury. Treatment of larvae with isotonic sorbitol did not affect the proportion of dextran-infiltrated larvae at any time point (∼17% for 10mpw and 0% for 20mpw), suggesting that perturbation of osmotic pressure gradients did not disrupt the time at which barrier function is reestablished (**Figure 4D**). However, treatment with blebbistatin significantly increased the proportion of dextran-infiltrated larvae during a dextran pulse between 7 and 10 mpw to ∼60% (**Figure 4D**, p=0.026 by Fisher’s exact test), consistent with previous reports that myosin contractility contributes to wound closure and sealing (Abreu-Blanco et al., 2012). These dextran infiltration experiments revealed that reestablishment of epidermal barrier function is promoted by tissue contractility, but not osmotic pressure gradients.

Although osmotic pressure and tissue contractility are not strictly required for external fluid entry, we hypothesized that they might affect the amount or pressure of the entering fluid and subsequently fissure formation. To test the role of osmotic pressure and contractility in forming fissures, we wounded larvae expressing mCherry in the basal layer in isotonic sorbitol or blebbistatin. Fissures appeared and propagated away from the wound in hypotonic conditions as described (**Figure 5A**, **Figure 1F**). In contrast, larvae wounded in isotonic sorbitol maintained an intact basal cell layer throughout the wound healing process (**Figure 5B**, **Video S5**). Larvae treated with blebbistatin developed fissures close to the wound, but these fissures failed to propagate further into the tissue (**Figure 5C**, **Video S6**).

**Figure 5.**
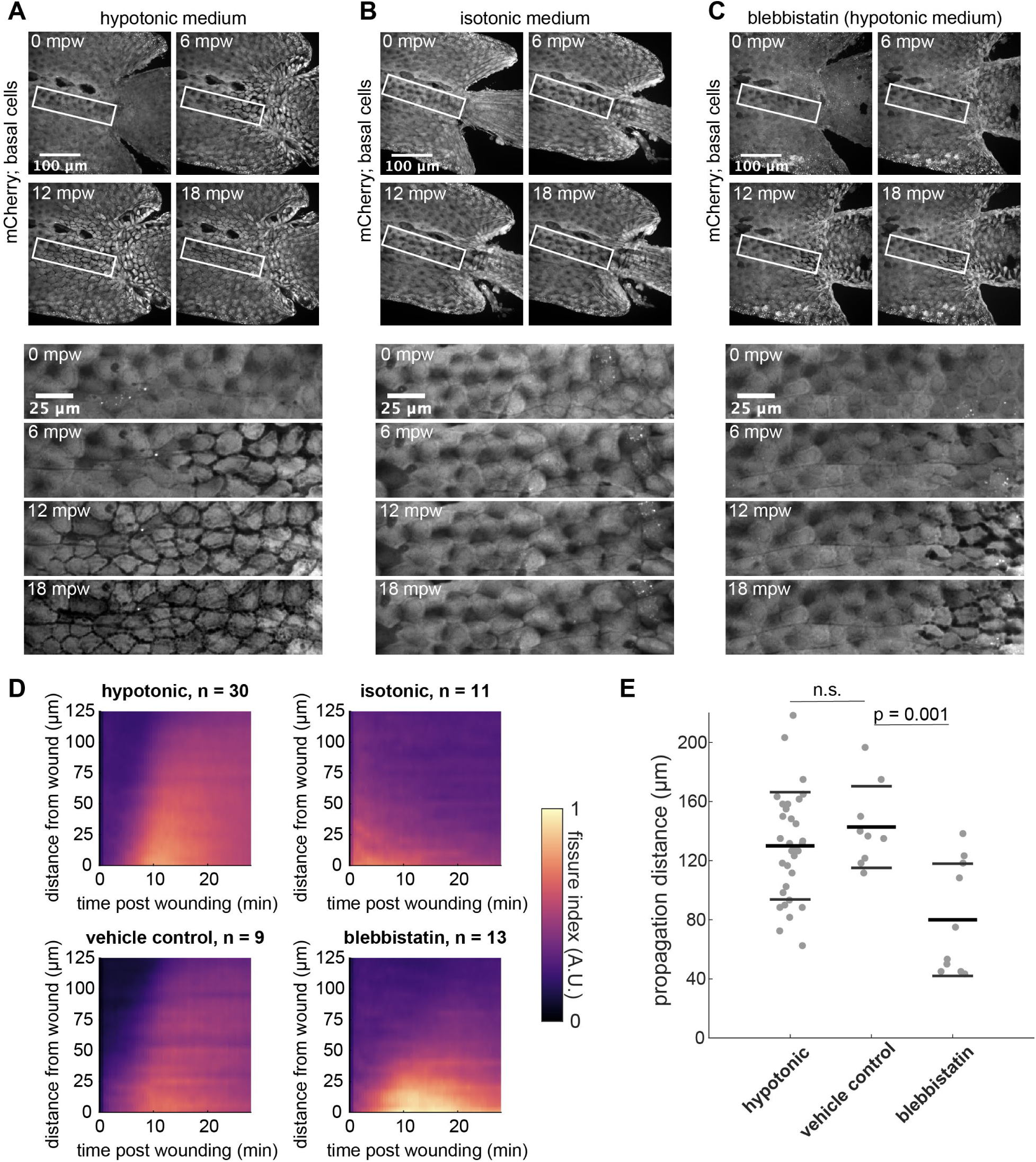
Osmotic pressure and tissue contractility promote fissuring. **(A)** *(Top)* Tailfin over time from a 3 dpf larva lacerated in E3 (hypotonic medium) and expressing mCherry in basal cells (*TgBAC(*Δ*Np63:Gal4); Tg(UAS:mCherry)*). Wound occurred 1-2 minutes earlier. *(Bottom)* Insets shown below the time-lapse series. **(B)** Larva wounded as in (A) in E3 supplemented with 270 mM sorbitol (isotonic medium). **(C)** Larva wounded as in (A) in E3 supplemented with 50 µM para-nitro blebbistatin. (A-C) are maximum-intensity Z-projections from spinning disk confocal images. **(D)** Kymograph indicating the fissure index over time at a given distance from the wound, averaged from larvae wounded in hypotonic E3 (*n* = 30) or E3 supplemented with 270 mM sorbitol (*n* = 11), 50 µM para-nitro blebbistatin (*n* = 13), or 0.1% DMSO as a vehicle control for blebbistatin (*n* = 9). The fissure index was analyzed as described in *Methods* and Figure 1. Data from larvae used in Figure 1F were incorporated into the average hypotonic fissure kymograph shown here (top left). **(E)** Quantified fissure dynamics for larvae that scored positively for fissure formation. Each dot is an individual larva (n ≥ 9). Thick and thin bars indicate the mean +/– 1 standard deviation for that condition. According to one-way ANOVA followed by Tukey’s test, vehicle control is not significantly differ from hypotonic (n.s., p > 0.05) and blebbistatin is significantly different from the vehicle control (p = 0.0010). Hypotonic data from Figure 1G is also plotted here for comparison.

To compare fissure propagation across conditions, we returned to our pipeline that quantifies local abundance of ridge-like structures as a fissure index. Averaging across larvae, fissure kymographs confirmed that the propagation of gaps observed in hypotonic media was absent in isotonic sorbitol (**Figure 5D**, top row). When fissure kymographs were scored for fissure formation (using blinded scoring as described in the Methods), all hypotonic-treated larvae (*n* = 30) and zero isotonic-treated larvae (*n* = 11) scored positively for fissure formation after wounding. Therefore, an osmotic pressure gradient between internal and external fluid is required to form large-scale fissures between basal cells.

Fissure kymographs averaged across blebbistatin-treated larvae showed a burst of fissuring with little propagation, whereas fissure propagation occurred normally in larvae treated with DMSO as a vehicle control (**Figure 5D**, bottom row). When these kymographs were scored, 100% of control larvae (*n* = 9) and 77% of blebbistatin-treated larvae (*n* = 13) formed fissures and were retained for further analysis. Manual measurement of the extent of linear propagation in the kymograph (performed by scorers blinded to the experimental condition) showed that fissures in blebbistatin-treated larvae propagated 80 µm on average, significantly less than the mean 143 µm propagation in control DMSO-treated larvae (**Figure 5E**, *p* = 0.0010 by Tukey’s test). These data demonstrate that tissue contractility promotes fissure propagation as well as wound closure. Although osmotic pressure gradients do not affect wound closure, they have a distinct role in forming fissures, perhaps by controlling the amount of external fluid entry.

### Fluid in fissures is cleared via macropinocytosis in the basal layer

So far, we have described the occurrence of fissures within the basal epidermal layer that form due to external fluid entering the wounded larva. In addition to fissures, we also noticed the presence of large intracellular vesicles in the basal cells near the wound site that were visible by all three imaging modalities—in light microscopy (**Figure 1D**, yellow arrowheads), histology (**Video S3**), and electron microscopy (**Figure 6C**). This led us to investigate the dynamics of endosomes during the wound healing process using an early endosome reporter: a tandem repeat of the FYVE domain (2xFYVE) tagged with GFP. The 2xFYVE domain binds specifically and with high affinity to phosphatidylinositol 3 phosphate (PI3P) that is found on the early endosomal membrane. In the absence of a suitable PI3P enriched endosomal membrane, this probe remains cytosolic (Gillooly et al., 2000). We therefore expressed EGFP-2x-FYVE in the basal layer (Rasmussen et al., 2015) to observe early endosome behavior during wound healing. We noted that at 10 mpw and onwards, there was a striking increase in the formation of large (>5 µm in diameter) 2xFYVE-labeled vesicles throughout the basal layer with a greater diversity in sizes relative to that observed pre-wounding and immediately post wounding (**Figure 6A**, **Video S7**, **Figure S4A**). Based on the presence of endosomes >0.5 µm in diameter, it is likely that these endosomes are derived from macropinocytosis—a highly versatile pathway characterized by endocytic vesicles greater than 0.5 µm in diameter (or ∼0.2 µm^2^ in cross-sectional area) (Hewlett et al., 1994). The large vesicle sizes attained in macropinocytosis can facilitate changes in a cell’s surface area. Thus, to observe the dynamics of macropinosomes during wound healing, we segmented the FYVE-labeled endosomes and pesudocoloured the population according to area into three categories: small (< 1 µm^2^ area, orange), medium (1-5 µm^2^ area, blue) and large (> 5 µm^2^ area, black). During segmentation, endosomes too close to be detected individually or containing non-uniform FYVE signaI were discarded (**Figure S4B**). Notably, segmentation provided only a rough estimate of the number and size of endosomes, as some structures were not identified or misidentified (**Figure S4B**).

**Figure 6.**
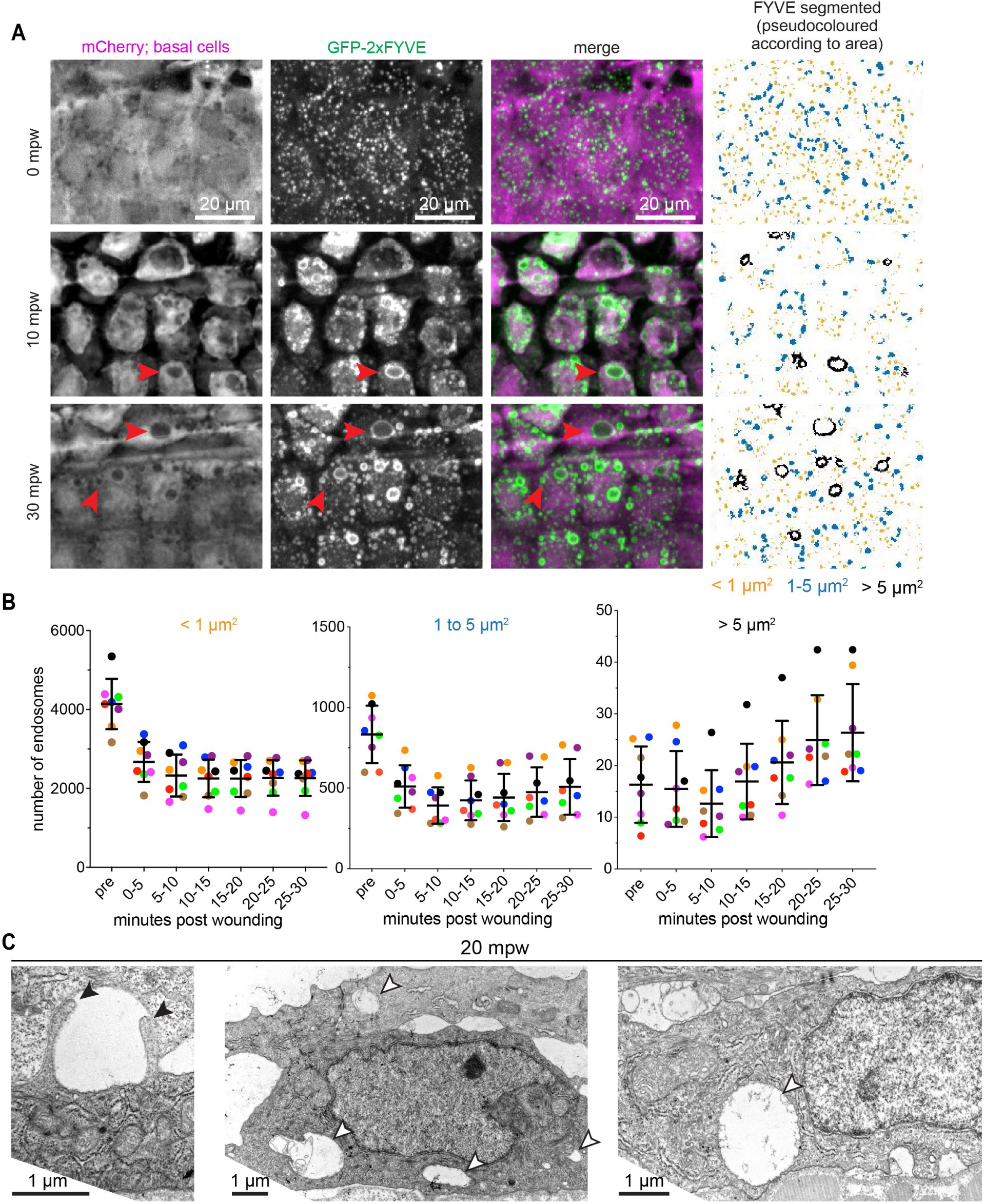
External fluid is cleared via macropinocytosis. **(A)** Maximum intensity projections from a timelapse following wounding of a 3 dpf larva expressing mCherry and GFP-2xFYVE in the basal cells that were wounded in the presence of E3 media (*TgBAC(*Δ*Np63:Gal4); Tg(UAS:mCherry); Tg(UAS:2xFYVE-GFP*)). First and second columns show the mCherry and GFP-FYVE channels respectively. Third column shows the merge. Fourth column shows segmentation of the GFP-FYVE. The segmented structures were divided into three categories and pseudocoloured (orange : < 1 µm^2^, blue: 1-5 µm^2^ and black: > 5 µm^2^). Red arrowheads show large endocytic structures. Full sized images are shown in Figure S4. **(B)** Graph shows quantification of mean endocytic structures over a 5 minute window as indicated for three different size categories (left: < 1 µm^2^, middle: 1-5 µm^2^ and right: > 5 µm^2^). Each fish is coloured uniquely (*n* = 8) and represented as dots. Lines represent mean +/- standard deviation across all the fishes. Note the difference in y-axis between the different plots. **(C)** Electron micrographs of basal cells in 3 dpf larvae treated with DMSO and wounded 20 minutes prior to fixation. White arrowheads highlight examples of large internal vesicles consistent with macropinosomes. Black arrowheads point out two protrusions from the apical side of a basal cell, possibly in the process of macropinocytosis.

We calculated mean endosome numbers over a 5 minute window to estimate population level changes over time post wounding (**Figure 6B**). Within 5 mpw, we observed a drop in numbers of both small and medium endosomes (**Figure 6B**), which corresponded to FYVE signal partially redistributed to the cytosol (**Figure 6A**, 10mpw). However, this drop in endosome number is probably overestimated due to tissue loss during the laceration. Thus, we focused on comparing the endosome number trends post wounding. Post wounding, the number of small and medium endosomes decreased over time relative to the first 5 minutes (**Figure 6B**, left and middle graph). In contrast, the number of large endosomes increased starting at 10 mpw (**Figure 6A**, red arrowheads and **Figure 6B**, right graph). Using electron microscopy we confirmed the presence of large vesicles within basal cells (**Figure 6C**). Intriguingly, we also observed an example of two protrusions extending into a fissure from the apical surface of a basal cell (**Figure 6C**, left panel), reminiscent of a cross-section through a macropinocytic cup (Veltman et al., 2016). Overall, our observations of large endosomes following fissuring suggest a scenario wherein fissures are generated by osmotic pressure and acto-myosin contractility and contain external fluid, and then this external excess fluid is subsequently cleared by the basal layer through macropinocytosis.

## Discussion

Taken together, our live imaging, electron microscopy, and histology results demonstrate that fluid influx during wound healing in larval zebrafish generates fissures between cells in the basal epidermal layer (**Figure 7**). Fluid influx is enhanced by the osmotic pressure gradient between the hypotonic external medium and isotonic interstitial fluid. As wound healing progresses, tissue contraction at the wound margin—likely mediated by the actomyosin purse string in the superficial cell layer—promotes the sealing of the wound, which increases hydrostatic pressure from the internalized fluid. This increase in pressure upon wound sealing leads to the formation of fissures via hydraulic fracturing or “fracking.” Fissures form at the lateral junctions of basal cells and along the interface between basal and superficial cells, and they propagate anteriorly from the wound margin for over 100 µm. Finally, the excess, externally derived fluid is internalized into large macropinocytic vesicles within basal cells, potentially in order to clear the fluid from the epidermis.

**Figure 7.**
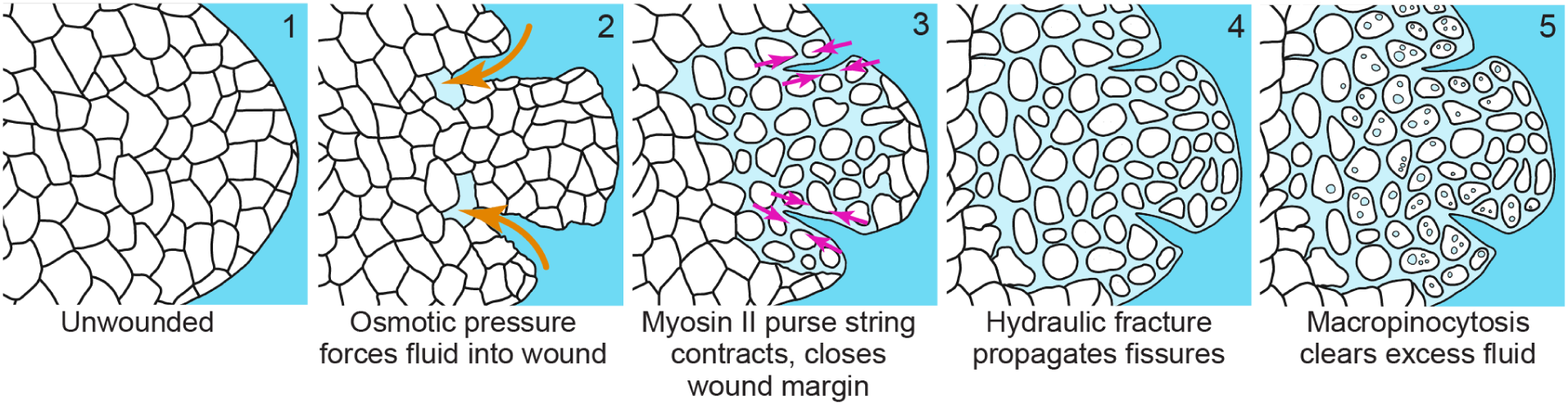
Model for wound-induced hydraulic fissuring. Overview of fissure formation during wound healing in zebrafish larval epidermis. (1) An unwounded tailfin showing an intact basal layer. (2) After wounding, osmotic pressure gradients drive external fluid through the open wound. (3) The myosin II purse string contracts along the wound margin, further forcing the external fluid anteriorly into the epithelial tissue. (4) The combination of the osmotic and cytoskeletal forces causes the hydraulic fracturing of the basal cells and fissure propagation anteriorly to the wound. (5) The excess fluid is subsequently cleared by the surrounding basal cells via macropinocytosis over longer timescales.

Osmotic gradients are required for fissuring, as fissures do not form when the external medium is isotonic with the internal medium (**Figure 5**). We believe that this is due to reduced fluid influx into the wound and not due to differences in the wound response *per se*. While isotonic solutions inhibit the wound response to some extent (Gault et al., 2014), isotonic sorbitol solutions such as those used in this work are still able to support a partial wound response involving directed cell migration of basal cells toward the wound and myosin II-dependent purse string contraction at the wound margin (Fuchigami et al., 2011; Kennard and Theriot, 2020). Therefore, we conclude the absence of fissuring that we have observed in isotonic medium is not due to a complete lack of wound response, but rather simply to the lack of a steep osmotic gradient.

In contrast to osmotic gradients, myosin II contractility is not strictly required for fissure formation, as fissures still occur in blebbistatin-treated larvae. However, myosin contractility is required for fissures to propagate anteriorly for long distances into the epidermis. We propose that this is due to the actomyosin purse string at the wound margin sealing the wound, which prevents fluid from flowing back out of the wound site. By closing off this path for fluid flow, tissue contraction builds up pressure in the epidermis, providing the conditions needed for hydraulic fracture and fissure formation. Consistent with this model, the typical time at which fissures first appear in the epidermis matches the time at which the wound becomes impermeable to dextran, around 7 mpw (**Figure 1G**, **Figure 4D**). Alternatively to the purse string model, cell-autonomous contractility within basal cells could promote fissuring deeper in the tissue. We favor the purse string model because it naturally suggests why fissures propagate anteriorly from the wound site, while the basal cell contractility model would require positing a spatio-temporal gradient in basal cell contractility to explain this propagation. Directly distinguishing between these two models will require the development of tissue-specific disruption of myosin contractility to specifically ablate purse string contractility separately from ablation of basal cell contractility.

Fluid accumulation due to swelling or flow of the extracellular matrix is an emerging mechanism for inducing morphogenetic tissue shape changes during development (Chugh et al., 2022; Munjal et al., 2021; Serna-Morales et al., 2022). The staining methods employed in our electron microscopy analysis are not diagnostic for extracellular matrix, so we cannot rule out the presence of extracellular matrix within fissures, which could contribute to fissuring due to hydrogel swelling effects.

Disruption of close cell-cell contact by hydraulic fracking was first described in cultured monolayers of mammalian epithelial cells growing on stretched hydrogels; when the stretch was released, excess water that had accumulated in the hydrogel was forced through the cell monolayer, leading to rapid junction disruption (Casares et al., 2015). In this setting, spatial propagation of hydraulically generated fissures was not observed, perhaps due to the small size of monolayers grown on micropatterns, or the experimental design, which led to the uniform release of water at all points in the hydrogel without lateral gradients in fluid flow. Hydraulic fracking has also been observed in the formation of the blastocoel cavity of the mouse embryo, where active ion transport generates an osmotic gradient that draws fluid into the embryo, leading to pressure-induced formation of lumens that gradually coalesce into the blastocoel (Dumortier et al., 2019). A similar process may underlie the formation of a single continuous lumen in the zebrafish gut (Alvers et al., 2014). Intriguingly, myosin II contractility promotes fissure closure in both epithelial monolayers and mouse embryos, while we have found that myosin II contractility instead promotes fissure formation and propagation in the context of wound healing. This discrepancy may be explained by a distinct role for myosin II contractility in generating permissive pressures for fissure formation in the wound setting.

The phenomenon we describe here of tissue fissuring driven by hydraulic fracking is quite distinct from tissue fracture, during which the epithelial layer rips apart leading to complete dissociation of formerly adjacent cells (Bonfanti et al., 2022). Such fractures have been investigated experimentally by pulling epithelial sheets beyond a critical strain (Harris et al., 2012), and fractures routinely occur in the course of normal movement in the simple multicellular organism *Trichoplax adhaerens* (Prakash et al., 2021). In contrast to tissue fracture, the wound-related hydraulic fissuring we report here features the retention of at least a subpopulation of cell-cell junctions, as seen at the ultrastructural level (**Figure 3B**). The tethers that connect basal cells during fissuring are reminiscent of the tethers cell use to retain contact with their substrate after partial retraction (Korkmazhan et al., 2022).

Our observation by both light and electron microscopy of dramatically large vesicles in the basal cells after wounding suggests that the accumulated external fluid is cleared by macropinocytosis. Macropinocytosis is a clathrin-independent process that generates large pleiomorphic vesicles, allowing a cell to respond quickly to changes in its environment in a way that enables the cell to change both its surface area and its net volume (Marsal et al., 2021). Macropinocytosis has been characterized most extensively in mammalian tissue culture cells in response to growth factors (Koivusalo et al., 2010; West et al., 1989), but there are also several cases where it seems to play a mechanical role. For example, constitutive macropinocytosis was reported in the outer epidermis of *Hydra vulgaris* that was downregulated by tissue stretch (Skokan et al., 2021). At a single cell level, immature dendritic cells generate macropinosomes that render them insensitive to hydraulic resistance, facilitating space exploration within the tissue (Moreau et al., 2019).

The regulated accumulation of interstitial fluid facilitates a variety of morphogenetic events in developmental contexts (Chugh et al., 2022). In addition to the formation of the blastocoel (Dumortier et al., 2019), relocation of interstitial fluid facilitates prechordal plate cell migration in zebrafish epiboly (Huljev et al., 2022), and interstitial fluid accumulation tunes the extent of zebrafish body axis elongation as part of a jamming transition in cell density (Mongera et al., 2018). Furthermore, osmotic swelling mediated by the accumulation of extracellular matrix components drives the formation of the semicircular canal of the zebrafish inner ear (Munjal et al., 2021). In all of these developmental processes, fluid flow is controlled internally, by changes in cell shapes or cell adhesion, or by generating osmotic pressure gradients through active transport or polarized synthesis of osmolytes. In contrast, the fluid that drives fissuring after wounding is externally derived, and fissuring represents a response to a rapid and extreme environmental change. Given such dramatic changes, the consistent and robust linear propagation of fissures anteriorly from the wound edge is all the more remarkable.

At this point it is not yet clear whether fissuring serves some function in facilitating other aspects of wound healing, as opposed to merely being an unavoidable, acute reaction by the animal to the wound and its consequent exposure to large volumes of dilute external fluid. However, it is tempting to draw a speculative parallel to the swelling that accompanies cutaneous wounds in humans (Lämmermann and Sixt, 2008; Scallan et al., 2010). Specifically, ”pitting edema” is defined as an indentation that remains behind after it is pressed, in contrast to healthy tissue, suggesting mechanical pilablity. Pitting edema is generated as part of the inflammatory response, during which blood vessel permeability increases, releasing fluid into nearby tissue and facilitating immune cell entry (Rutkowski et al., 2006). Although the source of additional fluid in inflammation is internal, rather than the external fluid that drives wound-associated fissuring in the fish, we speculate that the mechanisms deployed by the fish during wounding may have been refashioned in mammals to generate edema in response to inflammation and injury.

## Acknowledgments

We thank Jeff Rasmussen, Alvaro Sagasti and Holger Knaut for generously sharing fish lines. We are also grateful to Jeff Rasmussen, Rikki Garner and Maddy Hewitt for critical reading of the manuscript and thoughtful comments. We would like to thank LSB aquatics facility staff for animal care. We are also appreciative of Ksenia Pukhalskaya and Rita Chupalov for their help with Russian translation. Finally, we are grateful to members of the Theriot laboratory for numerous thought-provoking discussions. Electron microscopy was performed in the Biology Imaging Facility at the University of Washington. E.C.L was supported by an NSF GRFP (DGE-1656518). J.A.T acknowledges support from the Howard Hughes Medical Institute and the Washington Research Foundation.

## Author details

Conceptualization - A.S.K, J.A.T; Methodology - A.S.K., M.S., E.C.L., C.K.P., J.A.T.; Investigation - A.S.K., M.S., E.C.L., C.K.P.; Analysis - A.S.K., M.S., E.C.L; Illustration - C.K.P. Supervision and funding - J.A.T.; Writing - all authors. All authors read and approved the final manuscript.

## Methods

### Zebrafish husbandry

Zebrafish (TAB5 background) were maintained according to standard procedures. Experiments were approved by the University of Washington Institutional Animal Care and Use Committee (protocol 4427-01). Animals were maintained on a 14 hr light, 10 hr dark cycle at 28.5°C. Natural spawning was used for crosses, and embryos were raised at 28.5°C in 100 mm petri dishes in E3 without methylene blue (5 mM NaCl, 0.17 mM KCl, 0.33 mM CaCl2, 0.33 mM MgSO4) (“E3 medium,” 2008), with a maximum of 50 embryos per dish. All experiments were performed on larvae 72-90 hr post-fertilization.

### Transgenic zebrafish lines

The following previously generated transgenic zebrafish lines were used: *TgBAC(*Δ*Np63:Gal4)^la213^* (Rasmussen et al., 2015), *Tg(4xUAS:EGFP-2xFYVE)^la214^* (Rasmussen et al., 2015), *TgBAC(cdh1:cdh1-sfGFP)^sk95Tg^* (Yamaguchi et al., 2019), *Tg(UAS:LifeAct- EGFP)^mu271^* (Helker et al., 2013).

### Plasmid constructs and microinjection

To generate the *Tg(UAS:mCherry)* transgenic line used to visualize fissure in basal cells, the construct was cloned into Tol2kit zebrafish expression vectors using Gateway cloning (Kwan et al., 2007). Briefly, the *UAS:mCherry* plasmid was created using standard plasmids from the Tol2kit: p5E UAS, pME mCherry, p3E polyA, and recombined using Gateway cloning into a custom Tol2 destination vector (a generous gift from Darren Gilmour) with cry:mKate2 as a transgenic marker (Hartmann et al., 2020). To generate a stable transgenic line, wildtype embryos were injected at the 1- to 2-cell stage, into the cell (rather than the yolk). Plasmids were injected at a concentration of 20 ng/μl, with 40 ng/μl of Tol2 mRNA—the volume of these drops was not calibrated. Transgenic founders were identified by screening progeny for cry:mKate2 reporter expression, and subsequently confirmed by crossing with the *TgBAC(*Δ*Np63:Gal4)^la213^* reporter line.

### Preparation of larvae for imaging

Larvae were imaged at 3 days post-fertilization (3 dpf). Larvae were screened for transgenes in the morning prior to imaging. Larvae were prepared for imaging as previously described (Kennard et al., 2021; Kennard and Theriot, 2020). Briefly, larvae were anesthetized in E3 + Tricaine (E3 + 160 mg/l Tricaine (Sigma E10521) + 1.6 mM Tris pH 7) and mounted in 1.2-2% agarose (Invitrogen) in 35mm #1.5 glass-bottom dishes (CellVis D35-20-1.5N and D35C4-20-1.5N). Larvae were mounted with the *xy* (sagittal) plane parallel to the coverslip and excess agarose was trimmed from around the tail. E3 + Tricaine was the base for all experimental media. For isotonic treatment, larvae were pre-incubated for 10-60 minutes in E3 + Tricaine + 270 mM sorbitol (Sigma). For myosin inhibition, larvae were pre-incubated for 15 minutes in E3 + Tricaine + 50 µM *para*-nitro-blebbistatin (Fisher).

### Tissue laceration

Tissue laceration was performed as previously described (Kennard et al., 2021). Briefly, needles were formed from solid borosilicate glass rods (Sutter BR-100-10) by pulling in a Brown-Flaming type micropipette puller (Sutter P-87). This needle was used to impale the larvae just dorsal and ventral to the notochord and dragged at a 45 degree angle towards the posterior end of the tailfin to generate two cuts roughly following the orientation of actinotrichia (**Figure 1B**).

### Dextran entry experiments

For the fixed larvae experiments, typically 4-5 in parallel were anesthetized in E3 + Tricaine (E3 + 160 mg/l Tricaine (Sigma E10521) + 1.6 mM Tris pH 7) and mounted in 1.2-2% agarose (Invitrogen) in 35mm #1.5 glass-bottom dishes (CellVis D35-20-1.5N). Larvae were mounted with the *xy* (sagittal) plane parallel to the coverslip and excess agarose was trimmed from around the tail. Tricaine was added to all the experimental media. The chamber then was filled with one of three media: isotonic sorbitol, hypotonic E3, or blebbistatin (as described above). Tailfins of the larvae were lacerated (as described above) and were incubated (pulse) with 2mg/ml 10 kDa dextran [Alexa fluor 680 (D34680) & Alexa fluor 488 (D22910)] for 3 minutes following a 0, 7, or 17 minute wait time (chase). Next, the dextran was aspirated out of the chamber and larvae were washed twice with 2 mL of E3 + Tricaine. Larvae were washed twice with 60uL of 4% PFA (AA473479M, Fisher Sci Co) and were fixed with 30-60uL of 4% PFA for 20-25 minutes at room temperature. Larvae washed twice in E3 + tricaine and imaged in presence of E3 + Tricaine and were imaged (as described below) at room temperature.

For live experiments with dextran, larvae were mounted and prepared similar to the fixed larvae experiment. Tailfins of the larvae were lacerated in presence of 2 mg/ml Alexa Fluor 488 dextran and incubated for further 7 minutes. Larvae washed twice in E3 + tricaine and imaged in presence of E3 + Tricaine and were imaged (as described below) at room temperature.

### Light microscopy and image acquisition

Images were acquired on a Nikon Ti2 inverted microscope with a piezo-z stage (Applied Scientific Instruments PZ-2300-XY-FT), and attached to a Yokogawa CSU-W1 spinning-disk confocal with Borealis attachment (Andor), with illumination supplied by a laser launch (Vortran VersaLase) with 50 mW 488 nm and 50 mW 561 nm diode lasers (Vortran Stradus). Filters used included a 405/488/561/640/755 penta-band dichroic (Andor), a 488/561 dual-band emission filter (Chroma ZET488/561m) for rapid dual-color imaging, and single-band GFP, RFP, and far red filters for single-color imaging (Chroma 535/50m, 595/50m, and 700/75 respectively). An Apo 40x NA 1.25 water immersion objective was used for imaging (Nikon). Temperature control for live imaging was maintained with a resistive heating stage insert (Warner DH-40iL) with a temperature controller (Warner CL-100). Images were acquired on a back-thinned EMCCD camera (Andor DU888 iXon Ultra) in 16-bit mode without binning. Equipment was controlled using MicroManager v1.4.23 (Edelstein et al., 2010).

### Electron microscopy and histology

Larvae were anesthetized in a 35-mm plastic petri dish with E3 + Tricaine and lacerated under a dissection microscope as previously described (Kennard et al., 2021). At 2, 7, or 20 minutes post wounding, the tail was amputated just posterior to the yolk sac extension and the tail was immediately transferred with a flame-polished glass Pasteur pipette to 1.8 ml of freshly prepared fixation solution (1.5% Paraformaldehyde, 1.5% Glutaraldehyde, 0.1 M sodium cacodylate pH 7.4). Samples were fixed for 2-6 hours at room temperature, then washed 3 times for 10 minutes with gentle rotation in 0.1 M sodium cacodylate (all subsequent washes were performed with gentle rotation). Samples were stained with 1% (w/v) osmium tetroxide in 0.1 M sodium cacodylate for 2 hours at room temperature, washed 3 x 10 minutes with Type I water (MilliQ), then incubated in 1% (w/v) Uranyl acetate in water overnight at 4°C. Samples were dehydrated with successive 10 minute washes in an ethanol series (30, 40, 50, 60, 70, 80, 90, 95, 100, 100%) followed by two 10 minute washes in freshly opened acetone (Fisher). Samples were infiltrated for 1 h at room temperature in acetone with 33%(v/v) EMbed 812 resin, freshly prepared without accelerator (at a ratio of 22.6 g EMbed812 : 16.1 g DDSA : 9.8 g NMA, all from Electron Microscopy Sciences), followed by 1 h with 67% (v/v) resin in acetone, then 1 h in 100% resin, and then infiltrated overnight with fresh 100% resin. Samples were then incubated in resin with accelerator added (same resin with 2.5-3% (w/w) BDMA) for 2 h at room temperature. Samples were flat-embedded in a minimal amount of resin and baked overnight at 60 °C, and then hardened samples were cut out and re-embedded in fresh resin to orient the tails with the xz (coronal) plane parallel to the cutting plane. Re-embedded samples were baked at 60 °C for 48 hours.

Samples were trimmed with razor blades and sections were cut on a RMC PT-XL ultramicrotome. Wrinkles were removed by brief exposure to chloroform vapor. For electron microscopy, 60 nm (silver-grey) sections were cut using a diamond knife (Diatome Ultra 45°) and picked up on freshly plasma-treated copper slot grids with carbon-formvar coating (Electron Microscopy Sciences). For histology, ribbons of thick 500 nm sections were cut with a histo diamond knife (Diatome) and picked up onto ethanol cleaned pieces of #1.5 coverslip cut with a diamond pen to fit in the knife boat. To facilitate cutting serial sections, a dilute preparation of rubber cement in rubber cement solvent was sparingly applied to the top and bottom of the block.

Electron microscopy sections were post-stained with UranylLess (EMS 22409) for 1 minute and then 3% Lead Citrate (EMS 22410) for 6 minutes, and imaged on a Philips CM100 electron microscope at 80 kV with an Olympus Morada camera with iTEM software (Olympus).

Histology sections were stained for 30 s at 100 °C with azure B and basic fuchsin in sodium tetraborate as previously described (Morikawa et al., 2018) and mounted on glass slides in Permount (Thermo), which cured after 48 h at room temperature. After curing, excess Permount was removed from samples with a razor blade. Samples were imaged on a Zeiss Axioplan 2 upright microscope with brightfield imaging with a 100x NA 1.4 Plan Apo objective and an oil immersion condenser (NA 1.4). Images were acquired using a Nikon D3100 DSLR camera in RAW mode attached with a 2x relay lens. The camera was controlled using an intervalometer. To avoid blur, exposure times were set to roughly 1/500 s, and the camera mount was rotated with each acquisition to maximize the amount of tailfin present in the field of view. Vignetting was computationally removed by subtracting a background image and adding a constant offset to each frame. Images of serial sections were montaged and aligned using TrakEM2 (Cardona et al., 2012). Cells were reconstructed by manual tracing in napari, and reconstructions were visualized in ChimeraX (Goddard et al., 2018; Sofroniew et al., 2020).

### Surface projection and computational isolation of superficial and basal layers

To interpret dynamics within the basal layer, an alternative to maximum intensity projection was needed that would incorporate signal from just one side of the larva, and also separate the signal from basal and superficial levels of the epidermis. To this end we adapted the surface projection algorithm from the SurfCut ImageJ plugin (Erguvan et al., 2019), incorporating additional preprocessing steps and achieving increased processing speed by translating the algorithm to GPU programming. Briefly, raw data in the form of XY(C)ZT stacks were separated into individual XYZ stacks for each timepoint and channel, which were then background subtracted and flat-field corrected using fluorescent flat images collected as specified in (Model and Blank, 2006). The fluorescence of the flat was roughly matched to the intensity of the signal in the image, and then the corrected images were multiplied by a constant gain factor specific for each channel (20,000 for mCherry and 8000 for cdh1-sfGFP). Corrected stacks were then deconvolved with Huygens software (SVI) using constrained maximum-likelihood estimation and empirically measured point-spread functions specific for each channel. Following deconvolution, a surface projection of the mCherry signal from basal cells was generated using the SurfCut algorithm, rewritten to run on a GPU (Nvidia Titan Xp) using CLIJ2 (Haase et al., 2020). Unlike the original SurfCut algorithm, in which the threshold for detection of the surface was manually specified, in this custom implementation the signal was smoothed using a median filter and a Gaussian filter, and then thresholded using Otsu’s method. In parallel, the logarithm was applied to the smoothed image to create a second thresholded mask, and the union of the two masks was used; this step improved the thresholding of lower intensity regions.

The mask generated by SurfCut from the mCherry signal (restricted to the basal layer) was then used directly to isolate the E-cadherin signal from the basal layer. To isolate the superficial layer, the mask was translated towards the outside of the fish in the z direction by 6 µm.

### Fissure detection

Confocal timeseries of 3 dpf *TgBAC(*Δ*Np63:Gal4); Tg(UAS:mCherry)* larvae were processed for surface projection as described above. Following surface projection, movies were registered using previously-developed custom python code (Kennard and Theriot, 2020); registration transformations were computed on a 300 x 1024-pixel or 300 x 300-pixel (columns x rows) subimage furthest away from the wound. A straight line was drawn manually in Fiji to mark the position and orientation of the anterior-posterior axis of the embryo. Movies were spatially smoothed using a 5x5 median filter, and ridges were detected using Frangi’s algorithm implemented in MATLAB (Frangi et al., 1998; Kroon, 2010). A two-pass approach was used to select the Structure Sensitivity Factor (parameter *c* in Frangi’s original paper): the algorithm was first run on the final 10 frames of each movie, when fissures were certain to be prevalent in the image, with c chosen according to *c*^*^ = 0.5 × (*median*(*H*) + 10 × *MAD*(*H*)), where *H* is the Hessian norm of the image and *MAD* the median absolute deviation. The values of c^*^ from each of those 10 frames were averaged together and used as the parameter choice for the second pass through the entire movie, including regions in which little fissuring had occurred.

The Frangi ridge detector returns a value between 0 and 1 for each pixel that roughly corresponds to the probability that the pixel is part of a ridge. To generate a fissure kymograph, the anterior-posterior axis was manually defined, with the origin set at the wound site. The ridge detector’s output at each pixel was binned along the anterior-posterior axis in 5 or 10 µm increments by projecting each pixel’s coordinate onto the anterior-posterior axis line. Fissure largely propagated along the anterior-posterior axis, consistent with previous analysis of the propagation of cell movement in the tissue (Kennard and Theriot, 2020), justifying this approach. Ridge detector output for pixels in each 10 µm bin were averaged to generate a measurement at each time point at a given distance from the wound, which was visualized as a kymograph using custom MATLAB scripts (version 2018b, MathWorks).

### Quantification of fissure formation and propagation

Fissures kymographs were prepared from confocal timeseries of 3 dpf *TgBAC(*Δ*Np63:Gal4); Tg(UAS:mCherry)* larvae as described above. The fissure kymograph for each larva was evaluated using a custom Matlab script (version 2018b, MathWorks) by 3 independent researchers, who were blinded to the experimental condition. First, the kymograph image was converted into discrete points that marked the appearance of fissures. This conversion took advantage of the fact that fissures persisted to the end of the movie. Therefore, the fissure signal at a given distance from the wound over time approximated a step function. For each distance increment, a “transition point” was assigned at the time when the fissure index was half maximal (**Figure S2B***, top*).

Each larva was scored for fissure formation based on transition points overlaid on the original fissure kymograph. Kymographs were considered negative if the fissure index consistently decreased throughout the time-lapse and most transition points occurred at t=0 (**Figure S2B***, left*). Kymographs with fissures had a clear increase in fissure index at some point after wounding, indicated by transition points falling on a line with positive time-intercept and positive slope (**Figure S2B***, right*).

The distance, initiation time, and speed of fissure propagation were determined from each kymograph using a guided linear fit of the transition points. Each researcher marked the farthest distance of linear propagation and removed obvious outliers. A linear least-squares fit was performed on the remaining transition points within the linear region. Time was considered the response variable in order to properly fit short-distance, rapid propagation. Fits with slope below 0 or above 100 µm/min were excluded and the remaining parameters averaged between researchers (*n* ≥ 2).

### Bleaching correction

The signal from cdh1-sfGFP-expressing larvae faded quickly during imaging. To emphasize analysis of the spatial organization of E-cadherin rather than changes in signal levels (which could be due to photobleaching or protein degradation), images were bleach-corrected using the “Bleach Correction” function in Fiji, using the Simple Ratio Method with a Background level of 0.

### Endosome quantification

The wounded area was determined by hand-drawing the ROI around the wound and converting it to a mask. The mask was then multiplied to the images. The maximum Z-projection images were background corrected using a top-hat filter. The background corrected images were segmented using the threshold computed by the function ‘graythresh’ in MATLAB. After finding the endosome size using convex area function, endosomes were classified into three size groups: less than 1 µm^2^, 1-5 µm^2^ and greater than 5 µm^2^. Endosome number was averaged over the 5 minute interval.

### Code availability

Code used for image processing and analysis will be available on Gitlab (https://gitlab.com/theriot_lab/hydraulic-fissures-wound-healing) prior to publication.

