## Supplemental Materials for "Post-injury hydraulic fracturing drives fissure formation in the zebrafish basal epidermal cell layer"

*Contributed equally

### Supplemental Figures

#### Figure S1

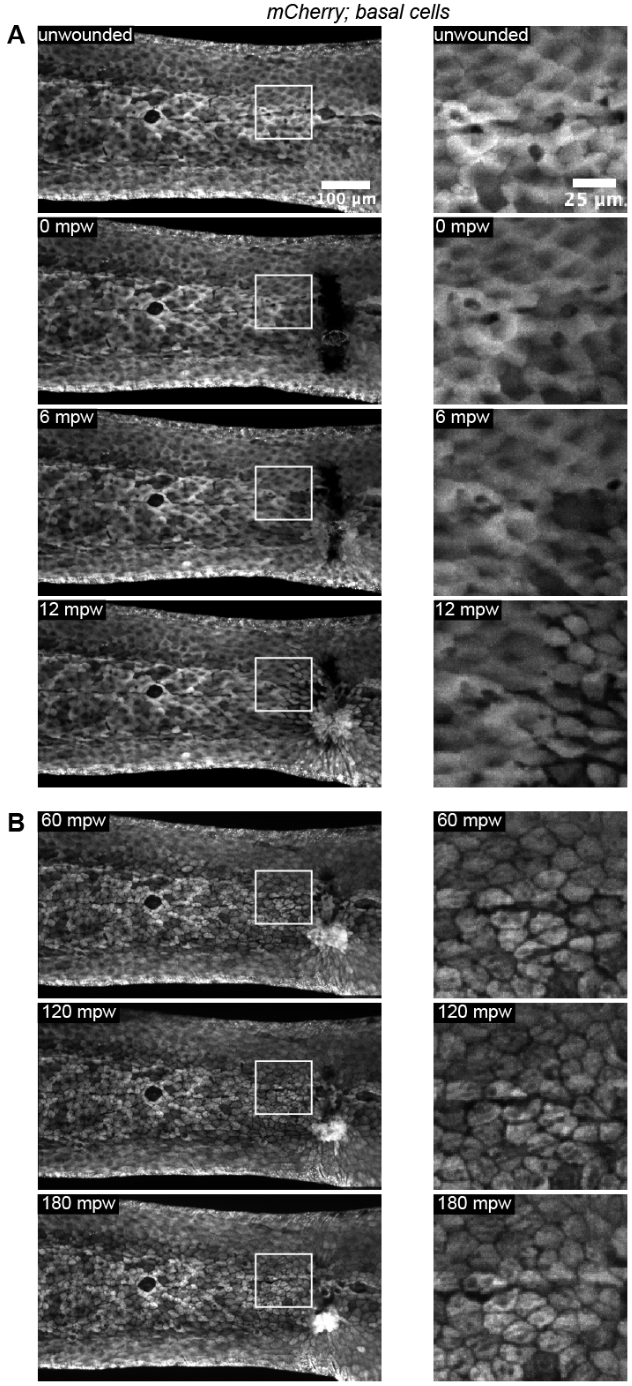

**Figure S1. Fissures persist for at least 3 hours after acute wounding.**

**(A)** Maximum intensity projections of 3 dpf larva expressing cytoplasmic mCherry in basal cells (*TgBAC(∆Np63:Gal4); Tg(UAS:mCherry)*) at different time points after laser wounding (405 nm, 120 mW diode laser, UGA-42 Firefly).

**(B)** Same larva as in (A), fissures persist at longer time points: 60, 120, and 180 mpw.

Figure S2

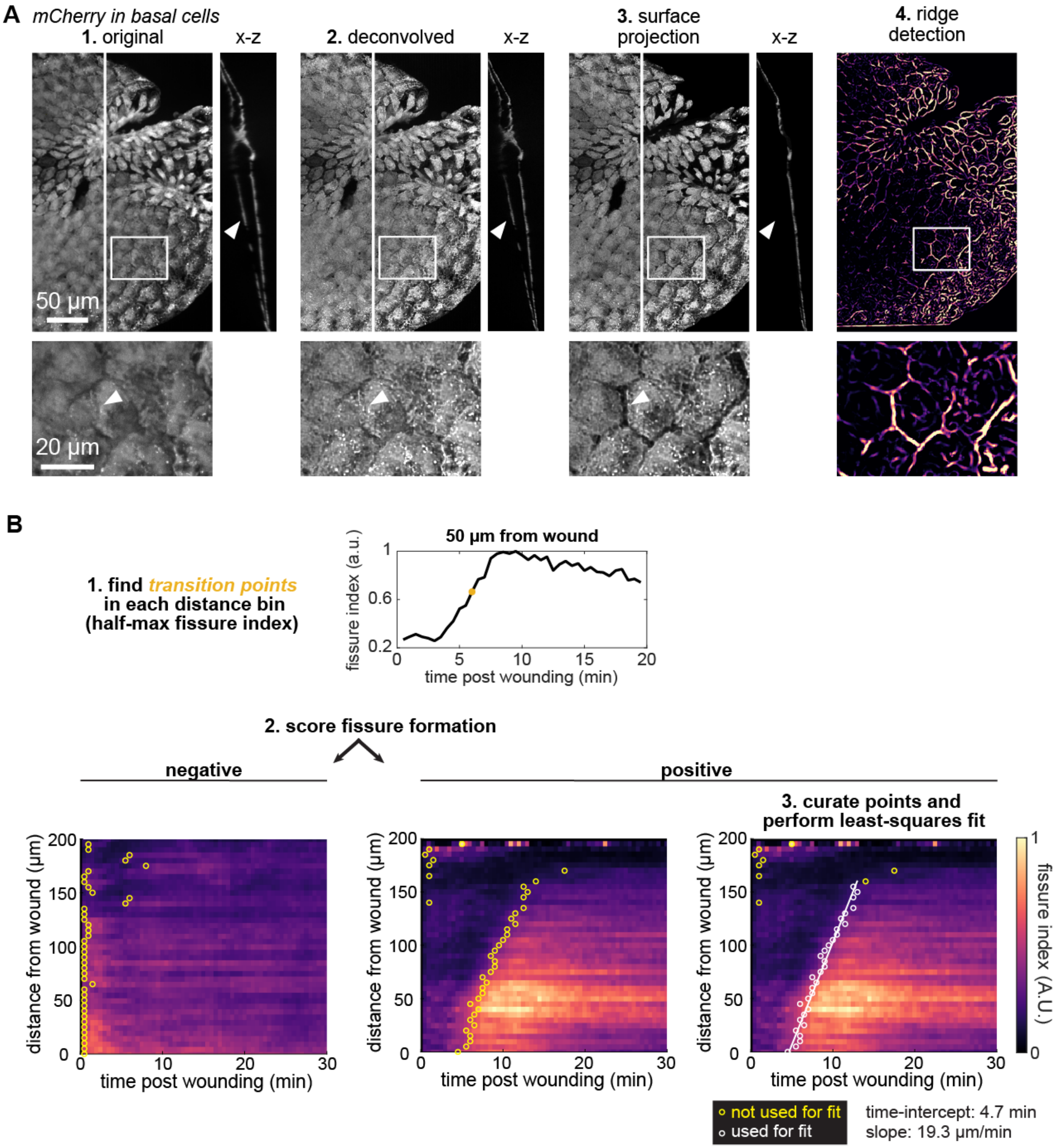


**Figure S2. Overview of quantification of fissure dynamics.**

**(A)** Illustration of the results of each step in the processing pipeline to identify fissures. Raw confocal stacks of mCherry expressed in basal cells were deconvolved, and then the epidermal surface closer to the objective was computationally isolated via surface projection, as seen in the x-z cross-sections. This step increased contrast of fissures because conventional maximum intensity projection would obscure fissures with signal originating from the opposite side of the larva. Following these preprocessing steps the 2D surface projection was filtered using a ridge detection algorithm that emphasized dark linear regions in the image. Example larva shown is the same as used in Figure 1E.

**(B)** To quantify fissure propagation dynamics from fissure kymographs, the spatiotemporal trend in fissure index was emphasized by identifying the transition point in each row of the kymograph (representing distance from the wound) where the fissure index attained 50% of the maximum value in that row. An example of the fissure index over time from a particular row of the kymograph is shown at the top of the panel, with the transition point marked in yellow. With transition points marked, kymographs were blinded and manually scored as depicting a fissuring event or not (see Methods), and for those kymographs judged to represent a fissuring event, the region of linear propagation was manually determined, facilitating the automatic extraction of a linear trend from the transition points with least-squares regression. Examples of negative and positive fissure kymographs are shown, and points that were marked for inclusion in regression are shown in white.

Figure S3

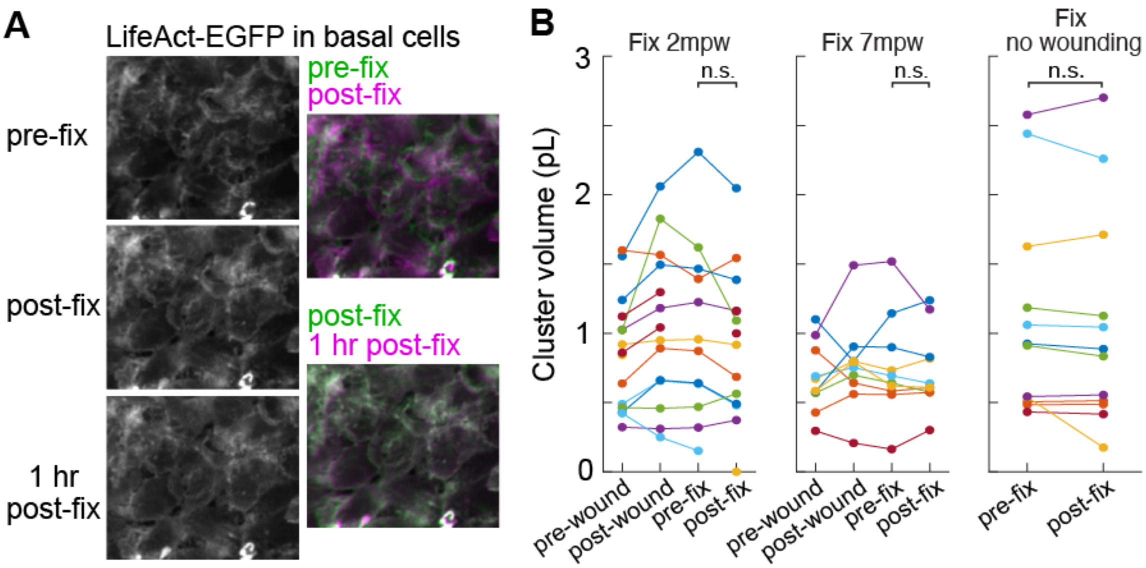


**Figure S3. Effect of fixation on tissue structure and cell volume**

**(A)** Maximum intensity projections from a 3dpf larva expressing LifeAct-EGFP in basal cells *(TgBAC(ΔNp63:Gal4); Tg(UAS:LifeAct-EGFP))*, before and after fixation with the buffer used for EM fixation. To better compare motion between the two frames, successive frames are overlaid in green and magenta at right. Note that roughly one minute elapses between the acquisition of the pre-fix and post-fix frames, so some of the movement between these frames is due to true tissue movement, rather than distortion during fixation.

**(B)** Cell volume was measured in mosaic clusters of cells expressing a fluorescent volume marker (mCherry) as described in (Kennard and Theriot, 2020), before wounding, after wounding, and before and after fixation. No significant trend was observed in cluster volume pre-fix vs. post-fix. n.s.: $p > 0.05$, paired two-sided $t$-test. For the panels left to right, n = 11, 10, 12 cell clusters from N = 4, 3, 5 larvae. Test was performed on all cells grouped together, regardless of larvae.

Figure S4

A

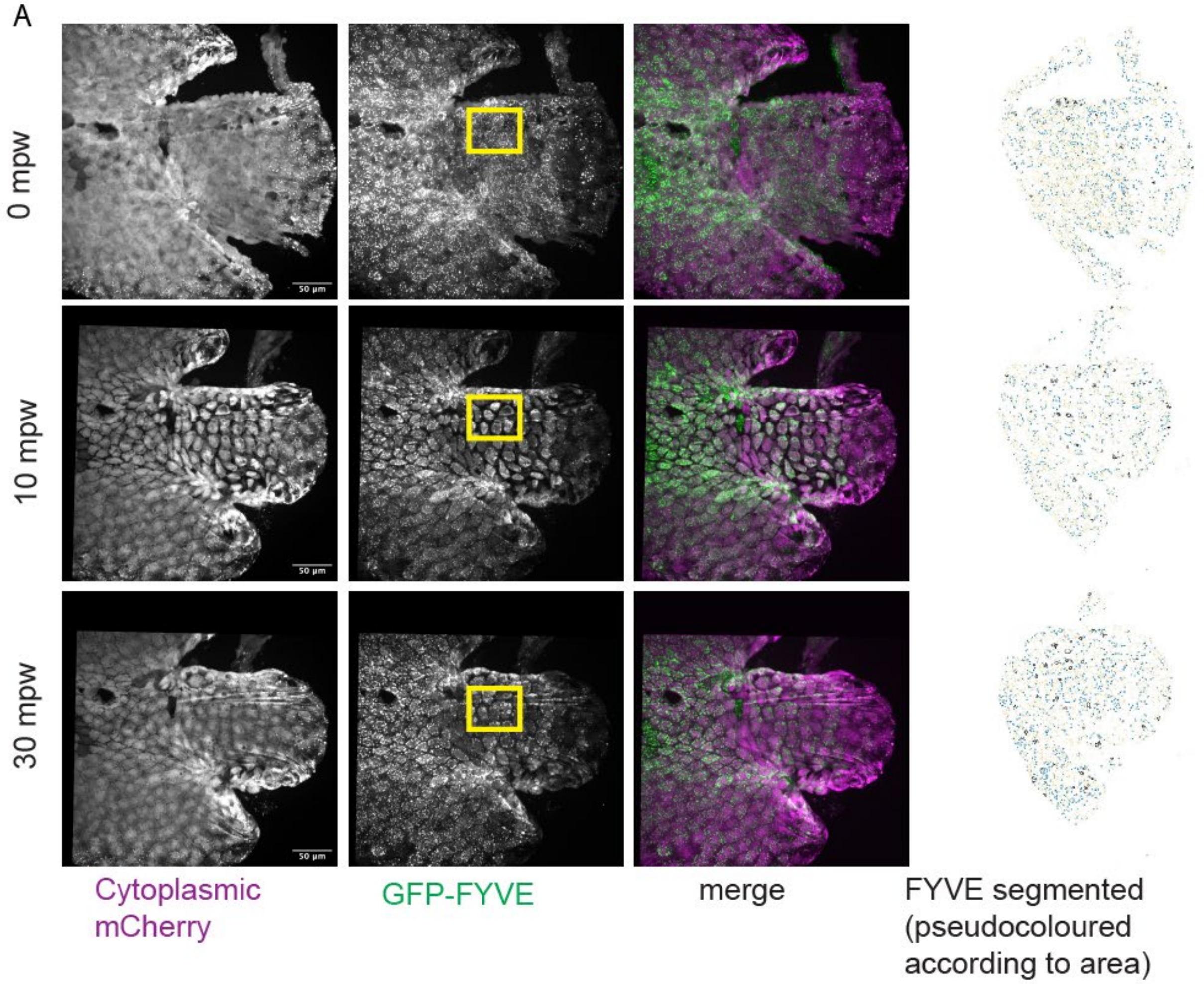

0 mpw
10 mpw
30 mpw
50 µm
Cytoplasmic mCherry
GFP-FYVE
merge
FYVE segmented (pseudocoloured according to area)


B

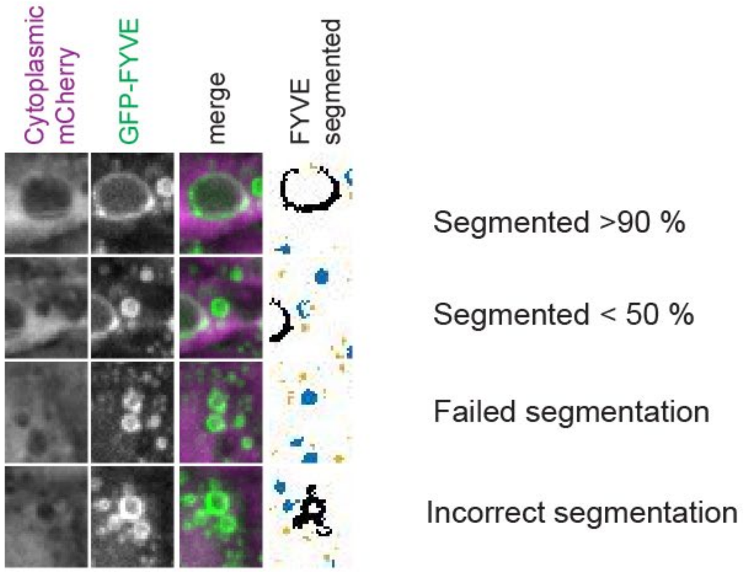

Cytoplasmic mCherry
GFP-FYVE
merge
FYVE segmented
Segmented >90 %
Segmented < 50 %
Failed segmentation
Incorrect segmentation

**Figure S4. Analysis of FYVE vesicle segmentation**

**(A)** Surface projection from a timelapse following wounding of a 3 dpf larva expressing mCherry and GFP-FYVE in the basal cells that were wounded in the presence of E3 media. First and second columns show the mCherry and GFP-FYVE channels respectively. Third column shows the merge. Fourth column shows segmentation of the GFP-FYVE. The segmented structures were divided into three categories and pseudocoloured (orange : $< 1\ \mu m^2$, blue: 1-5 $\mu m^2$ and black: $> 5\ \mu m^2$). Yellow box shows the inset shown in Figure 6A.

**(B)** insets from images shown in (A) depicting a successful segmentation of an endocytic structure (first row), partial segmentation of an endocytic structure (second row), failed segmentation of an endocytic structure (third row) and incorrect segmentation in which multiple vesicles were incorrectly merged into one structure (fourth row).

### Video Legends

**Video S1. Overview of wound-induced fissuring.**

Surface projection of 3 dpf larva expressing mCherry in basal epidermal cells, wounded by laceration. Fissures appear as dark gaps between cells, which propagate anteriorly from the wound site over the course of the video. mpw: minutes post-wound. Frames acquired every 30 seconds.

**Video S2. E-cadherin remains at cell-junctions throughout wound healing and fissuring.**

Wounded larva expressing mCherry in basal cells and zebrafish E-cadherin (Cdh1) in both basal and superficial cell layers. Shown are surface projections of mCherry in basal cells and E-cadherin computationally separated into signal emanating from basal or superficial cells. Video is of the same region as shown in Figure 3A. Note that the E-cadherin signal was bleach-corrected (see Methods) to emphasize differences in E-cadherin localization. To minimize photobleaching, frames were taken at non-uniform intervals: frames were taken every 30 seconds for 15 minutes, then every minute for 5 minutes, then every 2 minutes for 10 minutes, for a total of 30 minutes of acquisition. mpw: minutes post-wound.

**Video S3. Serial-section histology reveals interconnected fissures in epidermis.**

xz cross-sections of larva fixed at 20 mpw, embedded, sectioned, and stained for histology. Insets from this video are shown in Figure 3C. Video pans through 20 µm of sections along the y axis (dorsal-ventral), 0.5 µm per frame. The bi-layered epidermis is seen on the outermost edge of the larva, and the notochord, and muscles can be observed in the middle of the tissue. Collagen-rich appendages known as actinotrichia—precursors to the rays of the tail fin—are seen as dark purple elongated structures just beneath the epidermis. A number of cells are seen posterior to and peripherally connected to the rest of the tissue; these cells may have been injured during tissue laceration. Many large vesicles are observed within cells, especially near the wound, and gaps between basal cells and between the two layers of the epidermis appear to be contiguous with each other, consistent with the fissures observed with confocal microscopy.

**Video S4. External dextran enters via the open wound *in vivo* and in fixed larvae**

Movie shows surface projection timelapse following wounding of a 3 dpf larva expressing mCherry in the basal cells that were wounded in the presence of E3 media supplemented with 2mg/ml 10kDa TMR dextran. Top row shows mCherry in the basal layer, middle row shows dextran flowing through the fissures and the bottom rows shows the merge.

**Video S5. Isotonic external medium inhibits fissure formation.**

Maximum intensity projections of wounded 3 dpf larvae expressing mCherry in basal epidermal cells. Video on the left is from control larva in standard hypotonic E3 medium (same as in Video S6). Video on the right is from a larva treated with 270 mM (isotonic) sorbitol prior to and during imaging. Frames acquired every 30 seconds.

**Video S6. Myosin II contractility prompts fissure propagation.**

Maximum intensity projections of wounded 3 dpf larvae expressing mCherry in basal epidermal cells. Video on the left is from control larva in standard hypotonic E3 medium (same as in Video S5). Video on the right is from a larva treated with 50 μM para-nitro-blebbistatin, a myosin II inhibitor. Frames acquired every 30 seconds.

**Video S7. Basal cells accumulate large PI3P-vesicles after fissuring.**

Maximum intensity projections of wounded 3 dpf larva expressing 2xFYVE-GFP and mCherry in basal epidermal cells. FYVE is shown in green, mCherry is shown in magenta. Frames taken every minute.
